# Spatially organized cancer-associated fibroblast subtypes partition cutaneous carcinomas into immune-active and contracted, immune-repressed niches

**DOI:** 10.64898/2026.06.01.729186

**Authors:** Aschenbrenner Bertram, Forsthuber Agnes, Jessen Jakob, Krajic Natalia, Zheng Yimin, Hecke Alexander, Zhu Shaohua, Purkhauser Kim, Hasitzka Elisabeth, Schrenk Irene, Kholodniuk Daria, Silic Katarina, Soler Cardona Ana, Wollmann Eva, Tschandl Philipp, Gesslbauer Bernhard, Freystätter Christian, Radtke Christine, Petzelbauer Peter, Walko Gernot, Andre F. Rendeiro, Beate M. Lichtenberger

## Abstract

Basal cell carcinoma (BCC) and cutaneous squamous cell carcinoma (SCC) are the most common keratinocyte-derived malignancies, yet they differ markedly in invasiveness, metastatic potential, and immune contexture. Although cancer-associated fibroblasts (CAFs) are increasingly recognized as key regulators of tumor architecture and tumor immunity, the spatial organization of distinct CAF subtypes in cutaneous carcinomas and their functional relationship with immune cells remains incompletely understood. Using a 33-plex imaging mass cytometry (IMC) panel, we profiled 28 regions of interest (ROIs) from 17 human BCC and SCC specimens, encompassing more than 739,000 single cells, and integrated these data with RNA fluorescence in situ hybridization (RNA-FISH), immunohistochemistry (IHC), multiplex immunofluorescence, and in vitro functional assays. We identified four fibroblast populations, including immunomodulatory CAFs (iCAFs), matrix CAFs (mCAFs), myofibroblast-like CAFs (myoCAFs), and reticular fibroblasts (retFIBs), and found that aggressive tumor subtypes were characterized by increased stromal area, extracellular matrix deposition, and altered CAF composition. CAF composition differed most prominently across BCC subtypes, with nodular BCC enriched for mCAFs and infiltrative BCC showing increased myoCAF density, consistent with a shift toward a contractile stromal program. Spatial analyses revealed distinct CAF-immune niches: iCAFs localized to immune-cell-rich, inflamed niches enriched for activated and/or exhaustion-associated immune-cell marker programs, whereas myoCAFs occupied fibroblast-dense, immune-poor niches with globally reduced immune activation. mCAFs were preferentially associated with immune cell accumulation in the stroma and spatial immune compartmentalization, with limited immune cell presence within tumor nests. At the invasive front, CAF-immune coupling was highly subset-dependent, with iCAFs linked to antigen-experienced T-cell states and myoCAFs linked to immune exclusion. In vitro, patient-derived CAF cultures from myoCAF-rich biopsies showed enhanced collagen-gel contraction, with cultures enriched for MCAM^+^ CAFs displaying increased contractile capacity. Aggressive tumor variants displayed increased stromal nuclear YAP/TAZ, while complementary single-cell pathway analysis supported a mechanically remodeled stromal microenvironment in which mCAFs contribute ECM/matrix-remodeling programs and RGS5⁺/myoCAF-like populations show enhanced mechanotransduction-associated signaling, rather than a uniform CAF-wide increase in canonical YAP/TAZ transcriptional output. Together, these findings define spatially organized CAF programs in cutaneous carcinomas and identify myoCAF-rich stromal niches as a recurrent feature of aggressive, immune-repressed tumor architecture. These results nominate CAF composition as a biomarker of immune architecture and a potential determinant of therapeutic response.

## Introduction

Basal cell carcinoma (BCC) and cutaneous squamous cell carcinoma (SCC) are the two most common keratinocyte-derived malignancies, but they differ markedly in growth pattern, metastatic potential, and clinical risk. BCC usually follows a locally invasive course and only exceptionally metastasizes, whereas SCC carries a higher risk of destructive invasion, recurrence, and metastasis^1,2^. Within each entity, histopathological architecture further stratifies biological behavior. Nodular BCC typically forms circumscribed tumor nests, whereas infiltrative or sclerosing BCC grows as irregular strands within a remodeled, often desmoplastic stroma^3^. In SCC, well-differentiated tumors retain recognizable squamous maturation with keratinization and intercellular bridges, whereas poorly differentiated tumors show architectural disorganization, reduced or absent keratinization, increased atypia, and frequent mitotic activity^4^. These tumor-intrinsic and histological differences are accompanied by changes in the tumor microenvironment (TME), where fibroblasts, endothelial cells, immune cells, and epithelial cells interact to shape tissue architecture, invasion, and immune control^5^.

Cancer-associated fibroblasts constitute a major stromal compartment of the TME. Once viewed as a relatively uniform population, CAFs are now recognized as both highly heterogeneous and functionally plastic^6–8^. Across tumor types, several recurrent CAF states have been described, although nomenclature remains inconsistent between studies. Many frameworks distinguish primarily between myofibroblast-like CAFs (myoCAFs; α-smooth muscle actin (αSMA)-high, contractile) and inflammatory/immunomodulatory CAFs (iCAFs; cytokine– and chemokine-secreting), with additional frequently reported subsets including matrix CAFs (mCAFs; extracellular matrix (ECM)-producing) and antigen-presenting CAFs (apCAFs; MHC-II⁺). Functionally, these subsets influence immunity through complementary mechanisms. iCAFs can shape immune recruitment and polarization through cytokine and chemokine production, whereas mCAFs and myoCAFs with matrix-remodeling and contractile CAF programs alter tissue architecture, reinforce stromal barriers, and limit immune-cell access to tumor nests^9–17^. In skin cancer, the mCAF–iCAF continuum is particularly prominent. mCAFs predominate in nodular BCC and well-differentiated SCC preferentially localizing at the tumor-stroma border, whereas infiltrative BCC and poorly differentiated SCC show broad expansion of the CAF compartment with increased densities of both iCAFs and mCAFs^18^; however, mCAFs largely lose their preferential location at the tumor-stroma border.

Complementary single-cell and spatial analyses in skin cancer, lung cancer and head-and-neck squamous cell carcinoma show that CAF subtypes recur as spatially organized stromal-immune neighborhoods associated with immune exclusion and therapeutic responsiveness^14,19–21^. These studies also suggested lineage relationships among fibroblast subtypes, including links between tissue-resident fibroblasts, perivascular-like populations, and myoCAFs. Single-cell and spatial atlases across organs commonly place myoCAF programs as a relatively terminal, TGFβ-associated endpoint, typically reached along trajectories linking matrix/remodeling CAF states and inflammatory CAF states, with some studies additionally proposing contributions from tissue-resident fibroblasts and/or perivascular (pericyte) compartments^22^. In our previously published single cell RNA-sequencing dataset, trajectory structure and marker distributions support a model in which an RGS5⁺ perivascular-like state with partial myofibroblast-like features is most closely connected to tissue-resident (normal) fibroblasts, whereas iCAFs are positioned downstream of mCAFs^18^. The RGS5⁺ perivascular-like compartment likely represents a mixture of pericytes and myofibroblasts that cannot be distinguished at the transcriptional level^18^. Consistent with reports suggesting loss of vessel association during tumor remodeling, these cells appear to integrate into the tumor stroma and may contribute to ACTA2-high fibroblast pools^23^. Notably, Ganier et al. identify two molecularly distinct pericyte populations (RGS5⁺ and TAGLN⁺) in human skin and BCC, indicating that the perivascular compartment is heterogeneous and may overlap with ACTA2-associated stromal programs^24^.

Cords et al. performed large-scale IMC in >1,000 non-small cell lung cancer patients, defining 11 CAF phenotypes with distinct spatial distributions and demonstrating that CAF composition is an independent prognostic factor that stratifies patients into favorable vs unfavorable immune states and therapy response^25^. Other multiplex studies reinforce these principles: dedicated IMC panels map CAF niches in breast cancer^26^, deep IMC profiling in pancreatic ductal adenocarcinoma identifies CAF clusters enriched at tumor-stroma interfaces with specific immune associations^27^, and multiplex immunofluorescence in melanoma links CAF burden and CAF subtype markers to immune infiltration and outcomes^28^. Moreover, apCAFs were shown to localize near tertiary lymphoid structures and correlate with immunotherapy response^17^. Large-scale spatial transcriptomic atlases further reveal conserved CAF neighborhoods across multiple tumor types, suggesting generalizable stromal-immune circuits^29^. Together, these works indicate that CAF heterogeneity is not only quantitative but spatially organized into immunologically and prognostically relevant niches.

Recent advances in spatial proteomics, including IMC and related high-plex ion-based imaging platforms, enable in situ, high-dimensional mapping of multicellular tissue architecture at subcellular resolution^30–32^. These technologies have revealed tumor-specific immune niches, including TREM2⁺ myeloid clusters in BCC and stromal-immune circuits in SCC^33^, mapped immune landscapes in melanoma^34^, established practical frameworks for multiplex tissue analysis^35^, and identified CAF markers associated with aggressive BCC^36^.

In the present study, we use high-dimensional IMC to systematically chart CAF-immune interactions in BCC and SCC. We hypothesize that CAF subsets not only differ in abundance between tumor types but also organize into distinct immune niches that underpin their divergent clinical behaviors. By mapping these spatial stromal blueprints, we aim to provide mechanistic insights into immune marginalization in keratinocyte carcinomas and to identify CAF subtypes as potential biomarkers or therapeutic targets to enhance patient stratification and immunotherapy response.

## Results

### High-plex IMC atlas of keratinocyte carcinomas and their tumor microenvironment

We established a 33-plex imaging mass cytometry panel designed to capture epithelial, immune, endothelial, and fibroblast/mesenchymal lineages in keratinocyte carcinomas. Using this panel, we acquired IMC images from 4 nodular BCC (NOD BCC), 5 infiltrative/sclerosing BCC (INF BCC), 3 well-differentiated SCC (WD SCC), and 5 poorly differentiated SCC (PD SCC) samples. In total, 28 ROIs were ablated across 17 tumor samples, covering approximately 136 mm² of tissue, including 50 mm² tumor area and 86 mm² stromal area, and yielding more than 739,000 single cells after segmentation and quality control (**Fig. 1A,C**). The mean analyzed area was 8.0 mm² per sample. Histopathological classification was reviewed by a dermatopathologist. The BCC cohort included nodular BCC and conventional infiltrative/sclerosing BCC; importantly, micronodular BCCs were not included in the infiltrative BCC group analyzed here. This distinction is relevant because micronodular, infiltrative, and sclerosing BCC may be grouped together in simplified diagnostic schemes, whereas the stromal phenotype described in this study refers specifically to the infiltrative/sclerosing cases represented in our cohort^37^. Immune cells were identified using CD45, CD3, CD4, CD8, FOXP3, CD20, CD68, CD163, CD66b, MPO, and CD15. Epithelial cells were labeled with pan-KRT and E-cadherin, endothelial cells were identified by CD31, and fibroblast/mesenchymal markers included Vimentin, FAP, αSMA, TAGLN, COL11A1, MMP1, and IDO1. Of note, αSMA marked both myofibroblast-like cells and perivascular smooth muscle cells. CD56 highlighted NKT cells (CD56⁺CD3⁺) and a CD56⁺ non-immune population corresponding to nerve/muscle. Functional markers (PD-1, PD-L1, PD-L2, LAG-3, B7-H3, CD45RA, CD45RO, Granzyme B, Ki-67) profiled activation, inflammation, and exhaustion states. Nuclear and membrane stains (iridium intercalator and ISCK1/2/3) supported segmentation. Tumor, stroma, and vessels were annotated using a HALO-based classifier, and single-cell masks were generated for downstream analyses (**Fig. 1C**). All antibodies were titrated and validated; representative validation and segmentation are shown in **Supplementary Fig. 1**.

**Figure 1.**
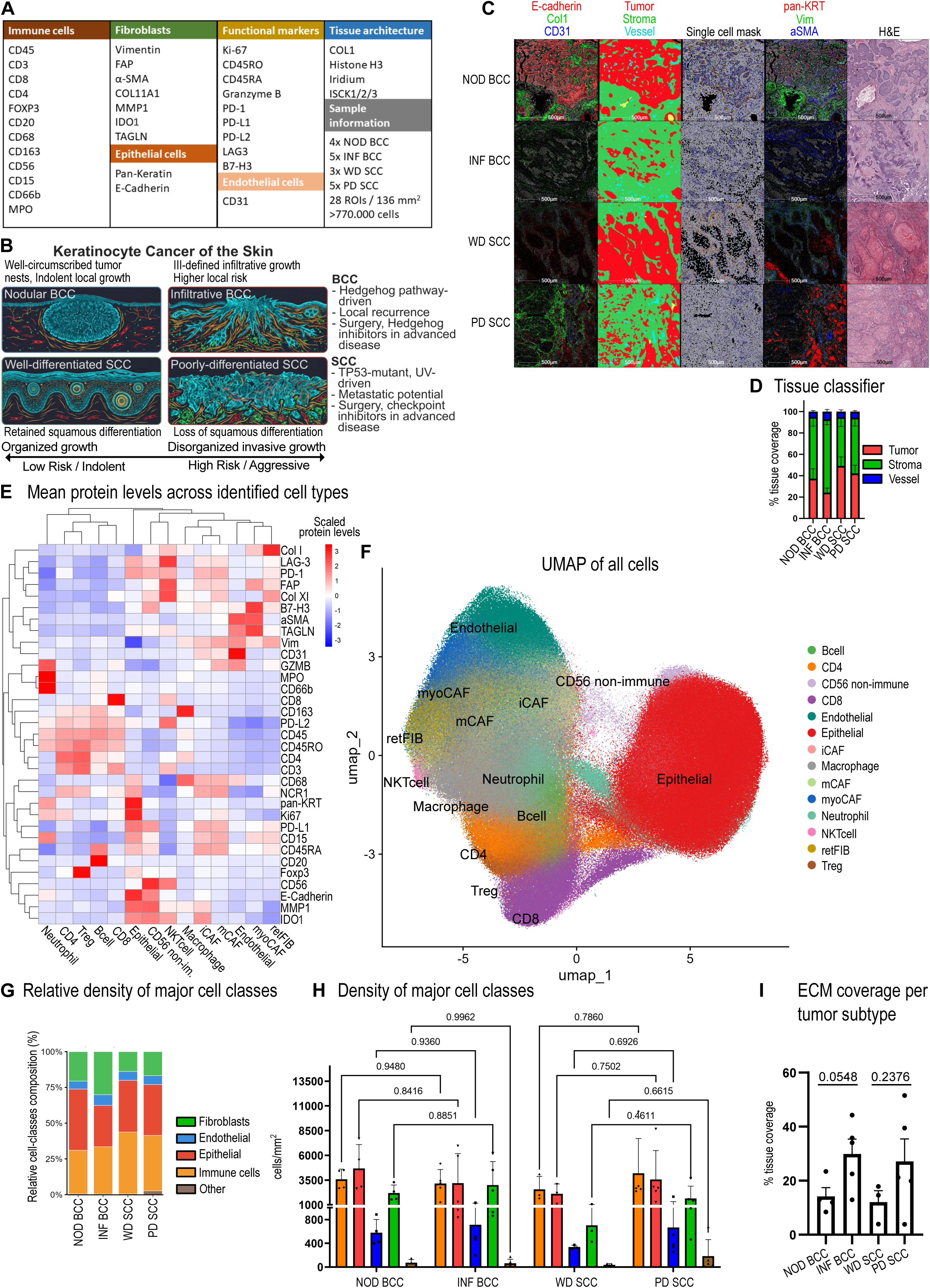
High-plex IMC maps tissue architecture, lineage composition and stromal expansion in keratinocyte carcinomas. **(A)** Overview of the 33-plex IMC antibody panel and sample cohort (17 tumors: 9 BCC, 8 SCC). **(B)** Schematic overview of the keratinocyte carcinoma subtypes analyzed in this study. NOD BCC is characterized by well-circumscribed tumor nests, whereas INF BCC shows irregular invasive strands within remodeled stroma. WD SCC retains squamous differentiation and keratinization, whereas PD SCC shows loss of differentiation, disorganized growth, and a more invasive phenotype. **(C)** Representative ROIs from BCC (nodular, NOD; infiltrative, INF) and SCC (well-differentiated, WD; poorly differentiated, PD). Columns (left→right): (1) E-Cadherin (red), COL1 (green), CD31 (blue); (2) tissue classifier (tumor = red, stroma = green, vessels = blue); (3) single-cell segmentation mask; (4) pan-KRT (red), vimentin (green), αSMA (blue); (5) H&E image. Images shown in pseudocolor. Scale bar, 500 µm. **(D)** Area fractions of tumor, stroma, and vessels per tumor subtype (mean ± SEM). **(E)** Heatmap of mean marker intensity per cluster (z-scored by marker; rows/columns hierarchically clustered). **(F)** UMAP of >700,000 single cells colored by cluster identity. **(G)** Relative density of major cell classes across tumor subtypes. Stacked bars show the mean relative contribution of epithelial cells, immune cells, fibroblasts, endothelial cells, and other cells per tumor subtype. **(H)** Absolute density of major cell classes across tumor subtypes. Bar plots show cell densities as cells/mm² for the same major cell classes. Points represent individual samples; bars indicate mean ± SEM. Exact p-values are indicated for selected planned comparisons. **(I)** Extracellular matrix burden from Masson’s trichrome whole-section images of the same tissue blocks as the IMC-analyzed sections (% tissue area classified as ECM).

Morphological comparison showed that INF BCC and PD SCC had larger stromal areas and vascular fractions, smaller tumor nests, and expanded tumor-stroma interfaces relative to NOD BCC and WD SCC (**Fig. 1C-D; Supplementary Fig. 2A**).

Unsupervised clustering using all markers, excluding Iridium and ISCK1/2/3, resolved the major epithelial, endothelial, and immune lineages, including B cells, CD4 and CD8 T cells, FOXP3⁺ Tregs, macrophages, neutrophils, and NKT cells, as well as nerve/muscle populations and two fibroblast-enriched clusters. To further define stromal heterogeneity, the two fibroblast-enriched clusters were merged and subclustered using the same marker set. This separated fibroblasts into myofibroblast-like CAFs (myoCAFs) and a broader non-myoCAF population. Subsequent re-clustering of the non-myoCAF population using selected fibroblast-associated markers, including MMP1, IDO1, COL11A1, and FAP, resolved three additional fibroblast populations: reticular fibroblasts (retFIBs), immunomodulatory CAFs (iCAFs), and matrix CAFs (mCAFs). In this step, MMP1/IDO1 and COL11A1 were the most informative markers for separating iCAFs and mCAFs, respectively, whereas FAP was retained as a broader CAF-associated marker and was not used as a subset-defining feature. Together with the αSMA⁺TAGLN⁺ myoCAF cluster identified during initial fibroblast clustering, this strategy defined four fibroblast populations: retFIBs, iCAFs, mCAFs, and myoCAFs (**Fig. 1E-F**).

retFIBs, representing healthy reticular fibroblasts, were classified based on both their low expression of CAF-associated markers and their spatial distribution, with the vast majority located at a distance from tumor cells, predominantly within the deep reticular dermis. Major cell-class composition was first visualized as relative fractions per tumor subtype (**Fig. 1G**). Absolute cell densities were then compared at the biological-sample level in cells/mm² for the planned within-entity comparisons NOD versus INF BCC and WD versus PD SCC (**Fig. 1H**). These density comparisons did not reach statistical significance but revealed consistent trends: INF BCC and PD SCC showed higher fibroblast densities than their less aggressive counterparts, while NOD BCC showed the highest relative immune-cell contribution. In SCC, WD SCC had lower absolute densities of both immune cells and fibroblasts than PD SCC; however, when expressed as relative fractions, immune cells comprised a larger share of WD SCC, consistent with a broader reduction across compartments rather than selective immune enrichment. Masson’s trichrome staining confirmed greater ECM deposition in INF BCC and PD SCC than in NOD BCC and WD SCC (**Fig. 1I; Supplementary Fig. 2F,G**).

### Conserved cell-type landscape across keratinocyte cancers, with CAF heterogeneity in BCC

We quantified relative lineage composition, extended immune-cell and CAF-subset distributions, and absolute cell densities across histological subtypes and tissue compartments (**Fig. 2A-C; Supplementary Fig. 3A-F**). Overall, epithelial cells constituted the largest compartment in each tumor group, with variable contributions from immune and stromal lineages (**Fig. 2A**). Extended composition plots further resolved immune-cell and CAF-subset distributions across tumor subtypes and individual samples (**Supplementary Fig. 3A-E**). In parallel, absolute density analysis quantified the abundance of all cell types across histological tumor subtypes within whole tissue, tumor, and stromal compartments (**Fig. 2B; Supplementary Fig. 3F**). Descriptive hierarchical ordering of subtype-level composition showed that NOD and INF BCC separated most clearly based on CAF-subset proportions, whereas this separation was not apparent when considering immune-cell composition alone (**Supplementary Fig. 3A,B**). This suggested that CAF subset composition, rather than broad immune-cell composition, represented the more prominent BCC subtype-associated difference.

**Figure 2.**
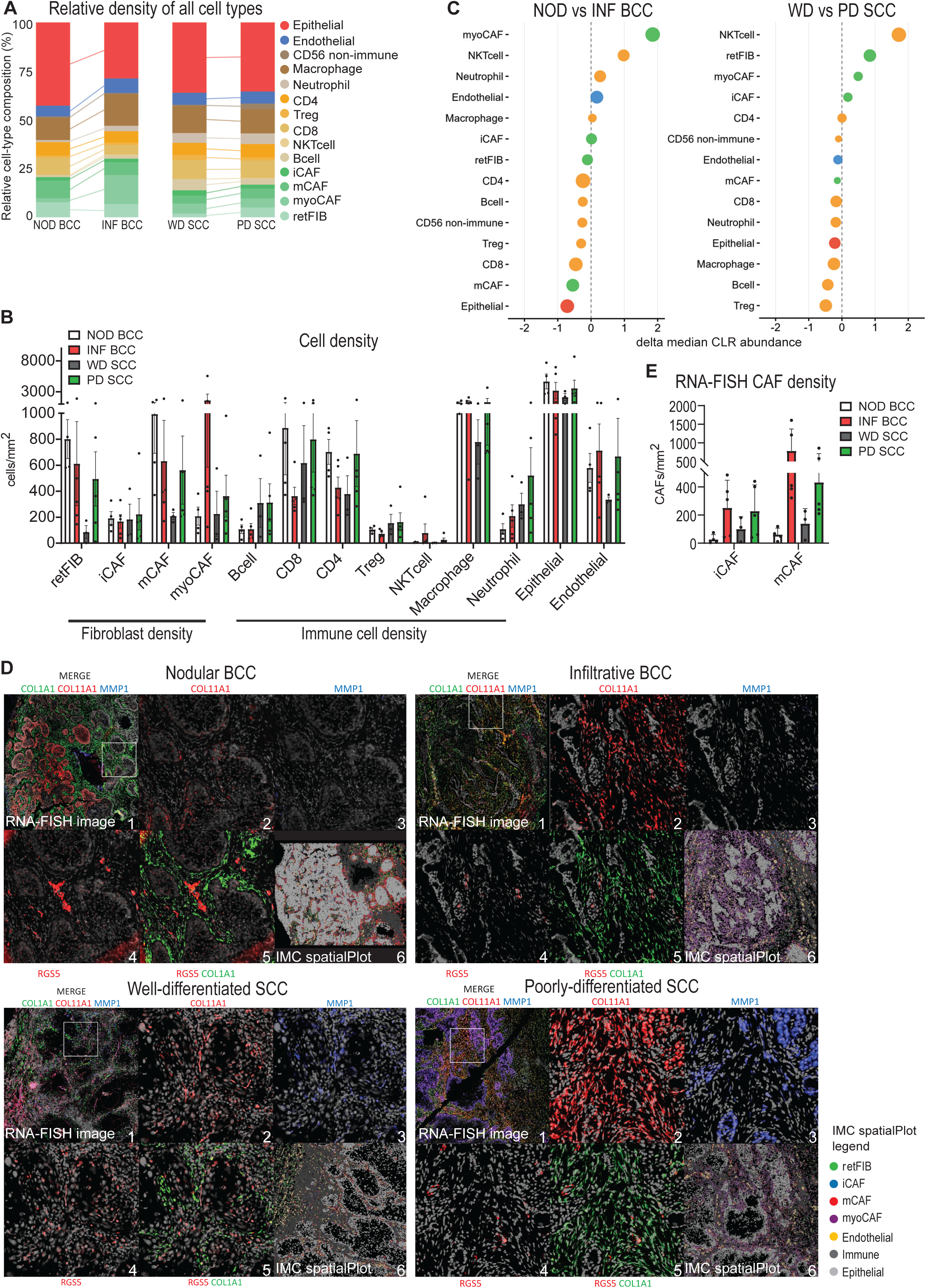
Sample-level composition and RNA-FISH validation reveal CAF-subtype shifts across keratinocyte carcinoma subtypes. **(A)** Relative lineage proportions among all cells per tumor type (%). **(B)** Mean cell densities per cell type across samples, grouped by tumor subtype (cells/mm²). **(C)** Sample-level compositional comparison of whole-tissue cell-type abundance between histological subtypes using centered log-ratio-transformed cell-type counts. ROIs were collapsed to biological samples before statistical testing. Ranked dot plots show the difference in median centered log-ratio abundance for each cell type. Positive values indicate higher abundance in INF BCC relative to NOD BCC or in PD SCC relative to WD SCC. Dot color indicates the major cell class, and dot size indicates −log10 nominal p-value from two-sided Wilcoxon rank-sum tests. **(D)** Representative RNAscope/RNA-FISH images with matched IMC scatterplots validating CAF subsets across all four tumor subtypes. 1: RNA-FISH overview at 1x (COL1A1 green, COL11A1 red, MMP1 blue), 2-5: zoom in to 20x magnification (2: COL11A1 red, 3: MMP1 blue, 4: RGS5 red, 5: RGS5 red & COL1A1 green), 6: IMC derived spatial scatterplot showing fibroblast subsets, endothelial cells, immune cells and epithelial cells in the color indicated by the legend. **(E)** Densities of iCAFs (COL1A1⁺ MMP1⁺) and mCAFs (COL1A1⁺ COL11A1⁺ MMP1⁻) determined by RNA-FISH of the same tissue blocks as the IMC-analyzed sections (cells/mm²).

To formally compare compositional shifts between histological subtypes while avoiding ROI-level pseudoreplication, ROIs were first collapsed to biological samples and centered log-ratio (CLR) analysis was performed on whole-tissue cell-type composition (Fig. 2C). Positive CLR differences indicate higher relative abundance in INF BCC compared with NOD BCC, or in PD SCC compared with WD SCC. Because NKT cells were rare in the dataset and retFIB abundance is sensitive to the amount of reticular dermis captured by each ROI, these ranked changes were interpreted cautiously. In BCC, the most biologically coherent compositional shift was the enrichment of myoCAFs in INF BCC, accompanied by reduced relative epithelial abundance, consistent with increased stromal area and the desmoplastic architecture of infiltrative/sclerosing BCC. Within the fibroblast compartment, NOD BCCs were dominated by mCAFs as previously shown^18^, whereas the infiltrative/sclerosing BCCs included in this cohort showed increased myoCAF representation, indicating a shift toward a more actomyosin– and ECM-remodeling fibroblast program in tumors with an invasive growth pattern^3^.

Because micronodular BCCs were not represented in our dataset, this myoCAF-rich stromal phenotype should not be generalized to all BCC subtypes that may be grouped diagnostically under broader “infiltrative” or “high-risk” categories. This interpretation aligns with evidence that CAFs in invasive carcinomas can display enhanced contractile and matrix-remodeling phenotypes, and that stromal remodeling and stiffening accompany invasive behavior^38–40^ (**Fig. 2B,C; Supplementary Fig. 3B,E,F**).

In SCC, CLR analysis indicated a broader compositional shift rather than a sharply defined CAF-state switch. CAF subtype composition was broadly similar between WD and PD SCC, while absolute fibroblast densities were lowest in WD SCC and endothelial cell densities were higher in the more aggressive histological subtypes (**Fig. 2B,C; Supplementary Fig. 3B,E,F**). Together, these data suggest that CAF-state remodeling is more pronounced in BCC, whereas SCC progression is associated with broader stromal and immune-context changes.

### Tumor aggressiveness is associated with a shift from nest-aligned mCAFs to stromal contractile myoCAFs in BCC

IMC-defined CAF subsets were compared with RNA-FISH on consecutive tissue sections to assess whether both modalities captured similar stromal organization at the tissue level. iCAFs defined by MMP1 and IDO1 protein abundance spatially overlapped with *MMP1* RNA-positive fibroblast-rich regions detected by RNA-FISH. mCAFs, defined by COL11A1⁺αSMA⁻ protein expression in IMC, aligned with *COL11A1*⁺*MMP1*⁻ fibroblast-rich stromal areas, particularly in NOD BCC. In contrast, IMC-defined myoCAFs, marked by αSMA and TAGLN protein expression, overlapped with *RGS5*⁺ myoCAF-like regions and with a subset of *COL11A1*⁺*MMP1*⁻ RNA-FISH-defined mCAF-like areas, particularly in stromal regions distal to the immediate tumor-stroma border (**Fig. 2D**).

RNA-FISH-stained sections were quantified using the workflow described in our previous publication^18^. Because *RGS5* expression in stromal cells was close to the detection threshold, *RGS5*⁺ stromal cells could not be robustly quantified. We therefore focused the RNA-FISH quantification on iCAFs and mCAFs. RNA-FISH-defined iCAFs (*COL1A1*⁺*MMP1*⁺) and mCAFs (*COL1A1*⁺*COL11A1*⁺*MMP1*⁻) were enriched in the more aggressive histological subtypes, particularly INF BCC and PD SCC (**Fig. 2E**)

To complement the visual comparison, we assessed sample-level concordance between IMC– and RNA-FISH-derived CAF densities using Spearman rank correlations (**Supplementary Fig. 4A,B**). Quantitative agreement between matched iCAF and mCAF definitions was limited, consistent with the fact that IMC and RNA-FISH capture different molecular layers and rely on partially divergent marker combinations. However, IMC-defined myoCAF density correlated positively with RNA-FISH-defined mCAF density, supporting the visual observation that protein-defined contractile CAFs overlap with a subset of transcriptionally defined *COL11A1*⁺ fibroblast states. Together, these data indicate that RNA-FISH and IMC converge at the tissue-organization level, while one-to-one quantitative agreement between CAF subsets remains limited by modality-specific RNA versus protein readouts and technical variation between consecutive sections.

### Protein-defined contractile programs distinguish mCAFs and myoCAFs

To directly link CAF transcriptional states to contractile marker expression *in situ*, we applied a single-section 5-plex assay combining RNA-FISH for *COL1A1, COL11A1, MMP1,* and *ACTA2* with αSMA protein immunostaining, yielding five simultaneous channels per tissue section (**Fig. 3A-D**). This analysis showed that *COL11A1*⁺ CAFs (mCAFs) were present across tumor types but exhibited tumor-dependent αSMA protein co-expression (**Fig. 3A,B,D**). In less aggressive lesions, a substantial fraction of *COL11A1*⁺ CAFs lacked detectable αSMA, consistent with a matrix-producing, low-contractility mCAF-like state rather than a myofibroblastic phenotype, and reflecting the separation of mCAFs and myoCAFs along contraction/myofibroblast and mural-associated programs described in our prior work^18^ (Fig. 3A,B,D). myoCAFs were enriched for actomyosin/contractility and mural markers, such as *RGS5* and *TAGLN*, whereas mCAFs exhibited low expression of these gene sets^18^. In contrast, in more aggressive tumors, *COL11A1*⁺ CAFs more frequently acquired αSMA protein expression, effectively shifting into a myoCAF-like phenotype and reducing the proportion of *COL11A1*⁺ CAFs that would be classified as “mCAF” when αSMA is used as a defining feature (**Fig. 3D**). Of note, we also found *COL11A1* expressed by nodular BCC tumor cells, however not in all tumor nests (**Fig. 3A**). By contrast, *MMP1*⁺ iCAFs exhibited little to no αSMA protein signal across different tumor types, indicating that iCAFs are largely αSMA-negative *in situ* and can be readily distinguished from contractile CAFs at the protein level. (**Fig. 3A,B,D**).

**Figure 3.**
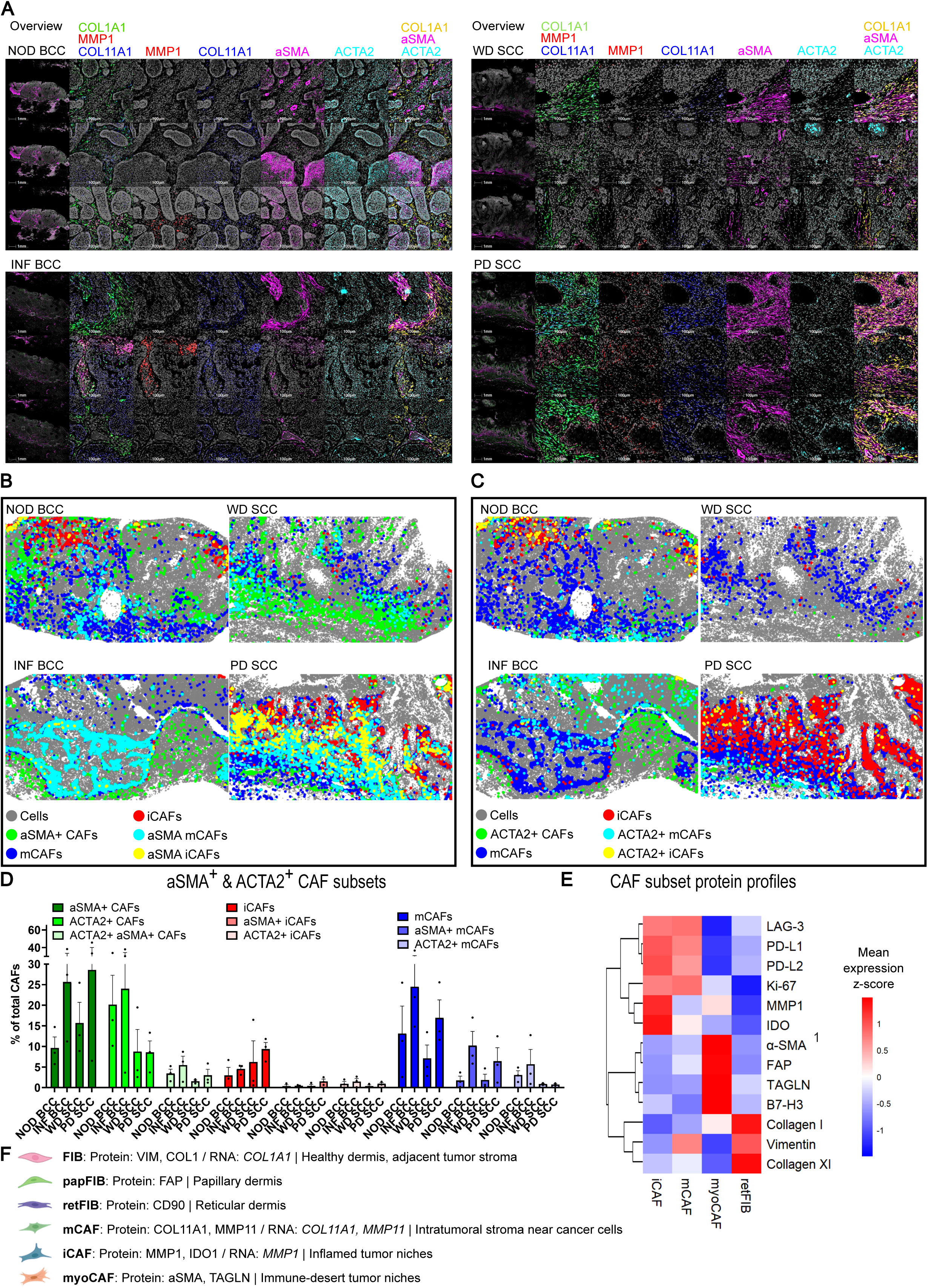
Integrated RNA-protein profiling separates matrix CAFs from contractile myoCAFs. **(A)** Representative RNA-FISH and antibody co-staining images of the investigated tumor subtypes. 4-plex RNA-FISH staining of COL1A1, MMP1, COL11A1, ACTA2, and αSMA on protein level. **(B)** Scatterplots validating CAF subsets positive for áSMA across all four tumor subtypes. **(C)** Scatterplots validating CAF subsets positive for ACTA2 across all four tumor subtypes. **(D)** Quantification of áSMA+ CAFs, ACTA2+ CAFs, ACTA2+ αSMA+ CAFs, iCAFs (MMP1+ CAFs), αSMA+ iCAFs, ACTA2+ iCAFs, mCAFs (COL11A1+ MMP1-CAFs), αSMA+ mCAFs and ACTA2+ mCAFs of 3 NOD BCCs, 3 INF BCCs, 3 WD SCCs and 3 PD SCCs, of the 4plex RNA-FISH co-stainings with αSMA antibody. **(E)** Mean IMC protein signal of CAF-associated markers across fibroblast subsets. Intensities were capped at the 99th percentile and z-scored per marker across subsets, indicating relative marker enrichment within the CAF compartment. **(F)** Graphical summary of fibroblasts subsets, their localization, RNA and protein markers in healthy skin and cutaneous carcinomas.

Across tumor types, αSMA protein and *ACTA2* RNA showed only limited concordance within CAFs (**Fig. 3D; Supplementary Fig. 4C,D**). Quantification of CAF subset composition showed that αSMA⁺ CAF fractions and *ACTA2*⁺ CAF fractions varied independently across samples, indicating that *ACTA2* RNA positivity is not a reliable proxy for αSMA-defined contractile CAFs (**Fig. 3D**). Consistently, pooled single-cell intensity analysis revealed only a weak association between αSMA and *ACTA2* levels (**Supplementary Fig. 4C**). Within-sample-centered comparisons further showed only a minimal increase in *ACTA2* RNA in αSMA⁺ CAFs compared with αSMA⁻ CAFs, with a median paired difference in sample-level median log1p(*ACTA2*) expression of approximately 0.061 (**Supplementary Fig. 4D**). Together, these analyses indicate that αSMA protein and *ACTA2* RNA capture only partially overlapping CAF states.

These findings also provide a framework for interpreting scRNA-seq CAF annotations derived from scRNA-seq. Since many single-cell atlases operationally define myoCAFs based on *ACTA2* transcript expression and often lack an explicit COL11A1⁺αSMA⁻ protein-defined mCAF state, COL11A1⁺ CAFs that acquire αSMA protein in aggressive tumors may be preferentially annotated as myoCAFs, whereas COL11A1⁺αSMA⁻ mCAFs may be underrepresented or subsumed within broader fibroblast categories^8,10,13,15^. This underscores the importance of integrating protein-level contractility markers with transcriptional programs when harmonizing CAF states across platforms.

IMC protein profiling revealed relative subset-associated marker patterns across CAF populations (**Fig. 3E**). The contractile myoCAF population separated during fibroblast clustering using the full marker set and was characterized by αSMA and TAGLN. By contrast, iCAFs, mCAFs and retFIBs were resolved within the non-myoCAF compartment, where MMP1, IDO1 and COL11A1 contributed to subtype annotation. These annotation-linked markers were therefore interpreted cautiously and not treated as independent functional discoveries. Beyond the defining markers, iCAFs and mCAFs showed comparatively higher PD-L1, PD-L2, LAG-3 and Ki-67 signals than myoCAFs and retFIBs, suggesting association with more immune-regulatory and proliferative stromal programs. myoCAFs showed the expected contractile profile, with high αSMA and TAGLN, together with higher FAP and B7-H3 signal within the CAF compartment and lower Ki-67 signal. retFIBs showed a structural matrix-associated profile, with higher Collagen I and Vimentin and comparatively low inflammatory and contractile marker signals. Collectively, these data indicate that CAF subsets differ not only by their annotation-defining markers but also by relative proliferative and immune-regulatory protein patterns.

Across samples, CAF marker-defined populations showed recurrent spatial organization when RNA-FISH and IMC protein maps were inspected together. In healthy skin (HS), fibroblasts are normally organized along the papillary-reticular dermal axis, with papillary fibroblasts located in the superficial dermis and reticular fibroblasts occupying deeper dermal regions. In the tumor ROIs analyzed here, however, papillary dermis was largely absent, and the non-CAF fibroblast population corresponded mainly to reticular fibroblasts located in deeper tumor-adjacent dermis. This interpretation is consistent with previous work defining human papillary and reticular fibroblast organization in skin^41^. Within tumors, CAF subsets showed distinct spatial tendencies: mCAF-like regions marked by *COL11A1*/*MMP11* RNA or COL11A1 protein were frequently located near tumor nests, iCAF-like regions marked by *MMP1* RNA or MMP1/IDO1 protein accumulated in inflamed stromal areas, and myoCAF-like regions marked by αSMA/TAGLN protein were most prominent in fibroblast-dense, immune-poor areas. These visual patterns support the interpretation that CAF subsets occupy distinct but partially overlapping tumor microenvironments rather than forming completely discrete compartments (**Fig. 3A,B,D,E; summarized in Fig. 3F**).

### myoCAFs are enriched in immune-desert tumors whereas iCAFs are enriched in inflamed tumors

Across tumor samples, we assessed coordinated variation in absolute cell densities using pairwise correlation matrices with hierarchical clustering (**Fig. 4A-B**). In the global analysis (**Fig. 4A**), immune lineages broadly co-varied, forming a positively correlated module that included CD4 T cells, CD8 T cells, Tregs, and macrophages. Neutrophils tended to be co-enriched with B cells. This pattern is consistent with tumors differing primarily in overall immune infiltration rather than exhibiting strictly mutually exclusive immune programs. Within this framework, iCAFs showed the most consistent positive associations, correlating with multiple immune cell types, including macrophages, CD4 T cells, and Tregs, suggesting that iCAF-rich tumors tend to coincide with immune-enriched microenvironments. mCAFs exhibited a similar but generally weaker pattern of positive correlations, most notably with macrophages and CD4 T cells, as well as with other stromal lineages (including retFIBs and endothelial cells), indicating that mCAF abundance tracks with immune cell-rich tumors, albeit less tightly than iCAFs. In contrast, myoCAFs displayed an opposing trend, showing predominantly negative correlations with many immune lineages and other stromal populations, positioning myoCAF-high tumors as compositionally distinct from immune-enriched samples (**Fig. 4A**). Consistent with this, density annotations and hierarchical clustering indicated that myoCAF-high tumors tended to group separately from tumors characterized by broad immune infiltration.

**Figure 4.**
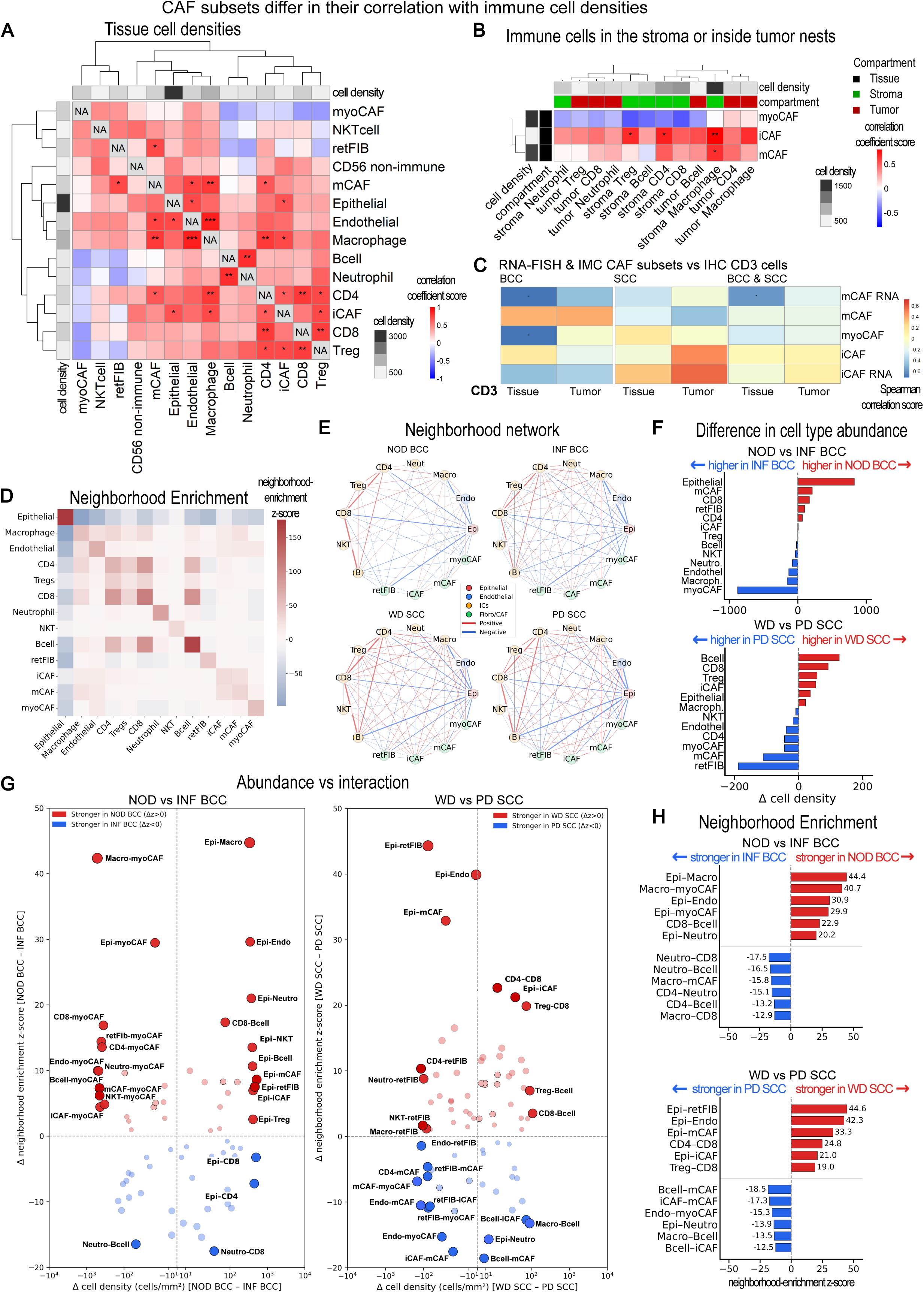
CAF subtypes segregate with distinct immune-density and neighborhood architectures. (**A**) Pearson correlation matrix (r) of tissue lineage densities across samples. Cell-type densities (cells/mm²) quantified by IMC in the tissue compartment were transformed as log(1 + density) prior to computing pairwise Pearson correlations across samples. Asterisks denote Benjamini-Hochberg (BH)-adjusted significance (*FDR < 0.05; **FDR < 0.01; ***FDR < 0.001). Side annotation indicates the mean density of each lineage across samples. (**B**) Pearson correlation matrix (r) of immune cell densities in stroma and tumor nests versus fibroblast subset densities in total tissue. IMC-derived fibroblast subset densities (iCAF, mCAF, myoCAF) in the tissue compartment were correlated against IMC-derived immune cell densities quantified separately in stroma and tumor compartments (cells/mm²), following log(1 + density) transformation, across samples. Correlations are shown for the CAF (tissue) × immune (stroma + tumor) block only. Asterisks denote BH-adjusted significance (*FDR < 0.05; **FDR < 0.01; ***FDR < 0.001), with multiple-testing correction applied separately for the stroma and tumor comparison families. Column annotations indicate immune compartment (stroma vs tumor) and mean density. (**C**) Spearman rank correlation (ñ) between CAF subset densities quantified by IMC and RNA-FISH (iCAF, mCAF, myoCAF, iCAFs.RNA, mCAFs.RNA) and CD3⁺ T-cell densities quantified by IHC in whole tissue and intra-tumoral nests. Cell densities (cells/mm²) were log10-transformed (log10[density + ε], ε = 0.01) prior to analysis. Heatmaps show correlation coefficients for all samples combined and stratified by tumor entity. Asterisks indicate nominal significance from two-sided Spearman correlation tests (*p < 0.05; **p < 0.01; ***p < 0.001; unadjusted p-values). (**D**) Mean neighborhood enrichment across all samples. Heatmap showing the mean Squidpy neighborhood-enrichment z-score for each cell-type pair across all samples. For each sample, ROI-level z-score matrices were first averaged to obtain one sample-level neighborhood-enrichment matrix, which was then averaged across samples. Positive values indicate that two cell types were observed as neighbors more frequently than expected by permutation, whereas negative values indicate neighborhood depletion. Color intensity reflects the magnitude of the enrichment or depletion score. (**E**) Neighborhood-enrichment networks by tumor subtype. Networks summarize subtype-specific neighborhood-enrichment patterns in NOD BCC, INF BCC, WD SCC, and PD SCC. Nodes represent cell types, and edges represent cell-type pairs with positive or negative neighborhood enrichment above the preset threshold (∣z∣≥1.96). Red edges indicate enriched neighborhoods, blue edges indicate depleted neighborhoods, and edge thickness scales with the absolute neighborhood-enrichment z-score. (**F**) Cell-type abundance differences between tumor subtypes. Bar plots show differences in mean sample-level cell density between tumor subtypes, calculated as Ädensity = NOD BCC − INF BCC or WD SCC − PD SCC. Red bars extending to the right indicate cell types with higher abundance in NOD BCC or WD SCC, whereas blue bars extending to the left indicate higher abundance in INF BCC or PD SCC. The dashed vertical line marks no difference in abundance. (**G**) Relationship between abundance changes and neighborhood-enrichment changes. Scatter plots compare changes in cell-type abundance with changes in neighborhood enrichment for NOD BCC versus INF BCC and WD SCC versus PD SCC. Each point represents one cell-type pair. The x-axis shows the difference in cell density between the paired cell types, calculated as Δdensity = A – B, and the y-axis shows the differential neighborhood-enrichment score, calculated as Δz = zA – zB. Red points indicate interactions stronger in NOD BCC or WD SCC, whereas blue points indicate interactions stronger in INF BCC or PD SCC. Displayed pairs were filtered for |Δz| ≥ 0.75, and the top-ranked pairs by absolute Δz are labeled. Dashed lines indicate no change in abundance or neighborhood enrichment. (**H**) Top altered cell-cell neighborhood enrichments. Ranked bar plots show the strongest subtype-associated changes in neighborhood enrichment between NOD and INF BCC or WD and PD SCC. Differential neighborhood-enrichment scores were calculated as Δz = zA − zB. Red bars indicate stronger enrichment in NOD BCC or WD SCC, whereas blue bars indicate stronger enrichment in INF BCC or PD SCC. The top altered cell-type pairs in each direction are shown.

We have previously shown that a high density of mCAFs at the tumor stroma border of BCC correlates with low numbers of T cells within tumor nests despite high T cell numbers in the adjacent stroma, indicating that mCAFs form a physical barrier preventing T cell infiltration^18^. To determine whether these relationships depended on immune-cell localization, we repeated the analysis after stratifying immune cell densities by compartment (stroma vs tumor nests) (**Fig. 4B**). This compartment-aware view reinforced the global patterns while adding spatial resolution. iCAF density was positively associated with stromal immune-cell densities, including CD4 T cells, Tregs, and macrophages, consistent with iCAFs co-occurring with immune infiltration. By comparison, mCAFs were less strongly aligned with stromal immune populations overall, but co-occurred particularly with stromal macrophages, while showing weaker or inverse relationships with immune densities within tumor nests. This pattern is consistent with a microenvironment in which immune accumulation is preferentially restricted to the stromal compartment and supports our previous findings that iCAFs represent a major cytokine/chemokine-producing stromal population, whereas mCAFs are associated with immune-cell exclusion from tumor nests^18^. Finally, myoCAFs again associated with an immune-low phenotype, showing negative correlations across most compartment-specific immune measures, consistent with myoCAF-rich tumors being comparatively immune-poor in both stroma and nests (**Fig. 4B**).

Overall, these analyses indicate that CAF subtypes segregate into distinct microenvironmental contexts: iCAFs mark immune– and vasculature-enriched tumors with immune-cell presence extending into tumor nests; mCAFs align more strongly with immune enrichment restricted to the tumor stroma; and myoCAFs characterize tumors with globally reduced immune-cell density.

To validate the CAF-immune density relationships inferred from IMC, we quantified CD3⁺ T cells by IHC on consecutive sections as an orthogonal pan-T cell readout (**Supplementary Fig. 2H**), extending our previous BCC-only analysis to include both BCC and SCC. INF BCCs showed lower whole-tissue CD3⁺ densities, while intratumor-nest CD3⁺ infiltration was comparable to NOD BCCs. In contrast, CD3⁺ densities in SCC were similar between WD and PD tumors. In BCC, RNA-FISH-defined mCAFs and IMC-defined myoCAFs were negatively associated with whole-tissue CD3⁺ density. In SCC, the inverse association between CD3⁺ density and RNA-FISH-defined mCAFs was weaker, and no corresponding association was observed for IMC-defined myoCAFs. Conversely, SCC showed a positive trend between iCAFs (defined by both RNA-FISH and IMC) and tumor-infiltrating CD3⁺ T cells (**Fig. 4C**).

### Tumor subtypes show spatial rewiring of CAF-immune cell neighborhoods

Permutation-based neighborhood enrichment analysis using Squidpy revealed recurrent spatial associations between major cell types across the dataset (**Fig. 4D-H**). Adaptive immune lineages preferentially localized near each other, with CD4 T cells, CD8 T cells, Tregs, and B cells showing mutual neighborhood enrichment. B cells also showed strong homotypic enrichment (**Fig. 4D**). Within the stromal compartment, iCAFs and mCAFs co-localized, and mCAFs were enriched near macrophages. In contrast, myoCAFs were depleted from neighborhoods containing most immune lineages, while retaining proximity to endothelial cells and other CAF subsets (**Fig. 4D**).

Subtype-specific neighborhood networks recapitulated these global patterns and revealed differences between tumor subtypes (**Fig. 4E**). Epithelial cells were mainly self-associated, consistent with the organization of tumor nests. Immune cells, especially CD4 and CD8 T cells, formed prominent interaction nodes across several tumor subtypes. NOD BCC showed the sparsest above-threshold neighborhood network, whereas INF BCC and PD SCC displayed more complex stromal and immune-associated neighborhood structures.

We next compared these neighborhood changes with subtype-associated differences in cell abundance. Cell-type density differences were calculated as NOD BCC – INF BCC or WD SCC – PD SCC (**Fig. 4F**). As expected, myoCAFs were increased in INF BCC, while PD SCC showed higher retFIB, mCAF, and myoCAF densities compared with WD SCC. In contrast, several immune cell populations, including B cells, CD8 T cells, Tregs, and iCAFs, had a higher density in WD SCC.

Differential neighborhood enrichment was then compared with these abundance changes (**Fig. 4G**). This showed that changes in cell density were not simply mirrored by changes in spatial association. For example, although myoCAFs were more abundant in INF BCC, many myoCAF-associated heterotypic neighborhoods were not stronger in INF BCC. Similarly, in SCC, PD tumors showed increased CAF densities, but only selected CAF-associated interactions, including iCAF-mCAF and endothelial-myoCAF neighborhoods, were stronger in PD SCC. Thus, neighborhood enrichment captured spatial organization that was not explained by abundance alone.

To summarize the strongest spatial changes, we ranked the top altered cell-cell neighborhood enrichments in each comparison (**Fig. 4H**). In BCC, NOD tumors showed stronger epithelial-macrophage, macrophage-myoCAF, epithelial-endothelial, epithelial-myoCAF, CD8-B cell, and epithelial-neutrophil neighborhoods. INF BCC showed stronger neutrophil-CD8, neutrophil-B cell, macrophage-mCAF, CD4-neutrophil, CD4-B cell, and macrophage-CD8 neighborhoods. In SCC, WD tumors showed stronger epithelial-retFIB, epithelial-endothelial, epithelial-mCAF, CD4-CD8, epithelial-iCAF, and Treg-CD8 neighborhoods. PD tumors showed stronger B cell-mCAF, iCAF-mCAF, endothelial-myoCAF, epithelial-neutrophil, macrophage-B cell, and B cell-iCAF neighborhoods.

Together, these analyses show that tumor subtypes differ not only in cell-type abundance, but also in the spatial organization of their epithelial, stromal, and immune compartments. Less aggressive or well-differentiated tumors retained stronger epithelial-stromal and epithelial-vascular neighborhood structures, whereas more aggressive subtypes showed altered immune-myeloid and immune-CAF-associated neighborhoods. This supports the concept that CAF-associated tumor progression involves spatial rewiring of the microenvironment rather than compositional change alone.

### CAF subsets preferentially occupy conserved spatial niches

The preceding neighborhood-enrichment analyses suggested that CAF subsets participate in distinct stromal–immune architectures, but these pairwise associations do not fully capture the multicellular composition surrounding individual fibroblasts. We therefore next focused directly on fibroblast-centered neighborhoods and asked whether marker-defined CAF subsets occupy recurring spatial contexts. Within a 100 µm radius, neighborhood composition differed across CAF subsets, indicating that CAF marker programs are embedded in distinct local microenvironments (**Fig. 5A**).

**Figure 5.**
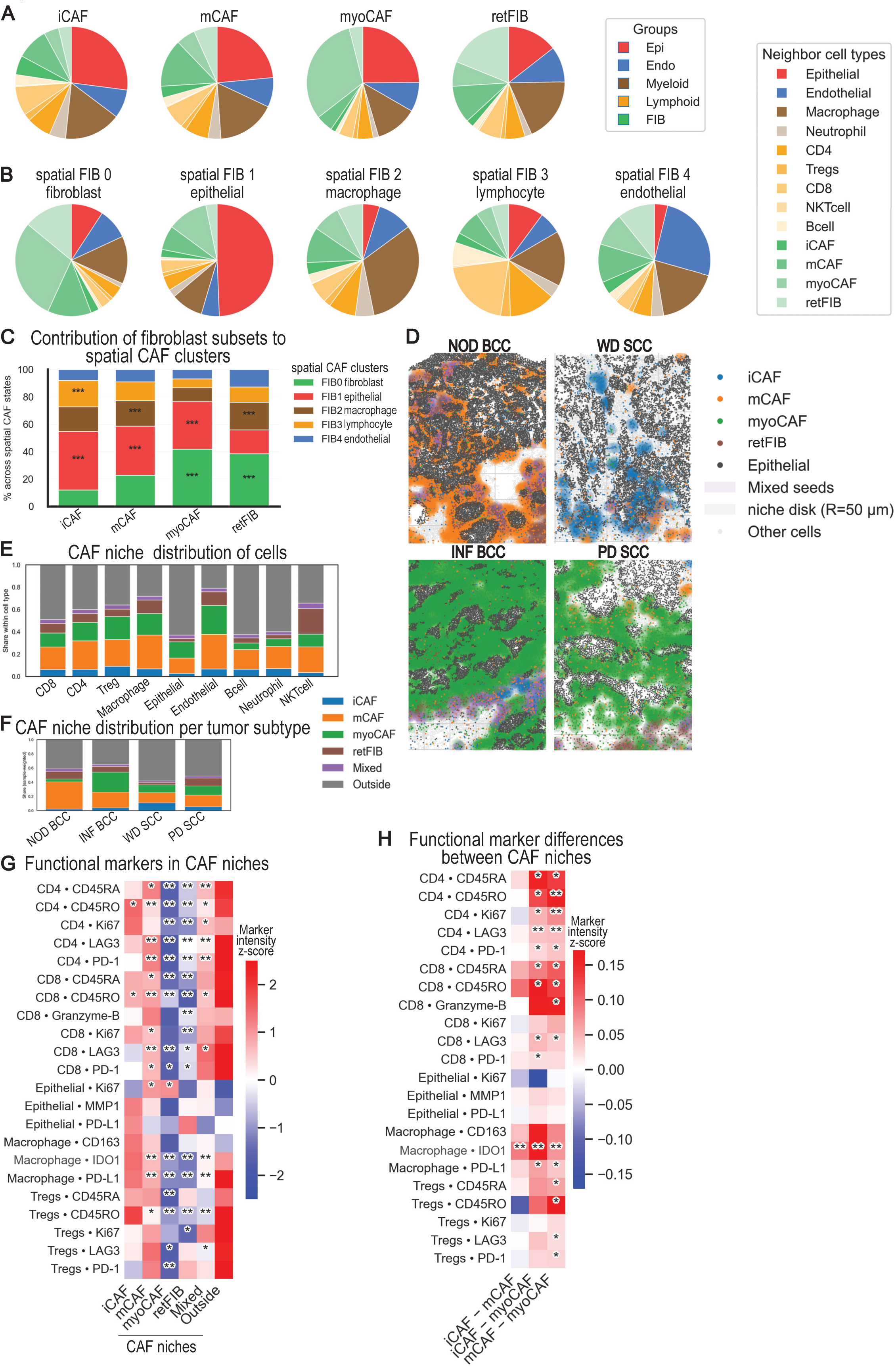
CAF-defined spatial niches shape immune cell activation and functional states. **(A)** Mean neighborhood composition for each CAF subset, calculated as the fraction of neighbors within a 100 µm radius around each fibroblast (centroid-to-centroid). Neighbor categories are grouped into epithelial, endothelial, myeloid (macrophage/neutrophil), lymphoid (CD4/Treg/CD8/NKT cell/B cell), and fibroblast classes; colors correspond to these groups and their subcategories. **(B)** Unsupervised identification of five spatial CAF neighborhood states (FIB0–FIB4) by non-negative matrix factorization (NMF) applied to fibroblast neighborhood composition (fibroblast neighbors collapsed into “FIB” for modeling). Pie charts show the mean neighborhood composition for fibroblasts assigned to each spatial state; state labels reflect the dominant adjacent compartment (e.g., epithelial-, macrophage-, lymphocyte-, endothelial-adjacent). **(C)** Spatial state distribution within each marker-defined CAF subtype (stacked bars; each bar sums to 100% within a marker subtype). Asterisks indicate marker–state pairs significantly enriched in the corresponding spatial state based on one-sided Fisher’s exact test with BH-FDR correction (enrichment defined relative to all other CAFs). **(D)** Representative scatterplots showing CAF-defined niches across the four tumor subtypes. **(E)** CAF niche distribution within major cell types: Stacked bar plot showing, for each major cell type, the fraction of cells assigned to each CAF niche class (iCAF, mCAF, myoCAF, retFIB, Mixed) or Outside. Cells were assigned exclusively to one niche class if located within at least one niche disk of that class. Outside denotes cells not inside any niche disk. Bars sum to 1 within each cell type. **(F)** CAF niche composition by tumor type: Stacked bar plot showing niche composition stratified by tumor type (NOD BCC, INF BCC, WD SCC, PD SCC). For each sample, the fraction of all cells assigned to each niche class (iCAF, mCAF, myoCAF, retFIB, Mixed, Outside) was computed, and tumor-type bars represent the sample-weighted mean across samples within each tumor type. Outside denotes cells not inside any niche disk. Bars sum to 1 per tumor type. **(G)** Pairwise Marker levels across CAF niches (row-wise z-score; CAF-scaled) with significance vs far-Outside: Heatmap summarizing marker expression across CAF niches for each cell type and marker. For each niche class, marker values were summarized per sample (median across cells within sample×class), then aggregated across samples (median). Heatmap values show a row-wise z-score computed using CAF niche columns only (iCAF/mCAF/myoCAF/retFIB/Mixed); the z-score is applied to all columns including Outside to aid interpretation of relative enrichment. Significance symbols indicate paired comparisons of each CAF niche class against the far-Outside baseline (cells at least 50 µm from any niche disk; see Methods) using paired two-sided Wilcoxon signed-rank tests with BH-FDR correction (* q<0.05, ** q<0.01, *** q<0.001, **** q<0.0001). Cells in the near-band between niche edge and far-Outside were excluded from these tests. **(H)** Pairwise CAF subtype differences in marker abundance (effect size heatmap). Heatmap showing pairwise differences between CAF-dominant niche classes (iCAF, mCAF, myoCAF) for each cell type–marker combination. Values represent effect sizes, calculated as the median of per-sample paired differences in transformed marker values (Δ = median[A − B]). Columns indicate the comparison direction (for example, iCAF − mCAF); red denotes higher values in the first class (A), and blue denotes higher values in the second class (B). Asterisks indicate significance based on paired two-sided Wilcoxon signed-rank tests across samples with Benjamini-Hochberg FDR correction (* q < 0.05, ** q < 0.01, *** q < 0.001, **** q < 0.0001). Only samples with non-missing values in both compared classes were included in each test.

To define these contexts in an unbiased but interpretable manner, we applied NMF to fibroblast-centered neighborhood composition and selected k = 5 to resolve the major local microenvironmental axes present in our cutaneous carcinoma dataset. This analysis was informed by pan-cancer spatial multi-omics work defining four conserved CAF spatial archetypes, including epithelial/tumor-adjacent, stromal/fibroblast-rich, myeloid-associated, and lymphoid-associated neighborhoods^29^. In addition, we allowed an endothelial/vascular-associated state to emerge, consistent with recent skin and barrier-tissue fibroblast atlases showing that fibroblast organization includes peri-vascular and other tissue-structured niches beyond canonical CAF classes^42,43^. Accordingly, the resulting spatial CAF states (FIB0–FIB4) corresponded to fibroblast-enriched, epithelial/cancer-enriched, myeloid/macrophage-enriched, lymphocyte-enriched, and endothelial/vascular-enriched neighborhoods, respectively (**Fig. 5B; Supplementary Fig. 4E**)^29^.

Mapping marker-defined CAF identities onto these spatial states showed that each marker class distributed across multiple neighborhood contexts, but with clear preferential enrichments (**Fig. 5C, Supplementary Fig. 4F).** iCAFs were significantly enriched in the epithelial/cancer-enriched state (FIB1) and the lymphocyte-enriched state (FIB3), indicating preferential localization within tumor–immune interface neighborhoods. mCAFs were significantly enriched in the epithelial/cancer-enriched state (FIB1) and the myeloid/macrophage-enriched state (FIB2), supporting an intermediate spatial phenotype bridging tumor-adjacent and innate immune-associated microenvironments. myoCAFs were significantly enriched in the fibroblast-enriched state (FIB0) and the epithelial/cancer-enriched state (FIB1), consistent with residence in fibroblast-dense stroma with extension toward tumor-proximal regions. retFIBs were significantly enriched in the fibroblast-enriched state (FIB0) and the myeloid/macrophage-enriched state (FIB2), indicating preferential association with stromal and innate immune-skewed niches rather than a single exclusive compartment.

Together, these results support a model in which CAF marker programs do not map one-to-one onto spatial microenvironments. Instead, marker-defined CAF subsets occupy a spectrum of conserved neighborhood states, with significant enrichments highlighting biases toward specific epithelial-, immune-, stroma-, or vessel-associated contexts.

### iCAF niches define inflammatory, checkpoint-high immune-cell states, whereas myoCAF niches are immune-repressed

To investigate whether CAF-defined neighborhoods associate with immune and epithelial functional states, we defined CAF niches by placing 50-µm-radius disks around CAF “seed” cells (iCAF, mCAF, myoCAF, retFIB) and assigning each non-fibroblast cell to one of six categories based on localization: iCAF, mCAF, myoCAF, retFIB, mixed-CAF (reflecting CAF neighborhoods without a dominant local subtype), or outside any CAF niche (Outside) (**Fig. 5D-H, Supplementary Fig. 4G**). This approach allowed us to move beyond CAF abundance and ask whether distinct CAF subsets are linked to specific functional states in neighboring immune and epithelial cells.

Spatial maps and summary bar graphs (**Fig. 5D,F**) revealed marked tumor-subtype differences in CAF niche engagement and niche composition. BCC samples showed a larger CAF niche-associated fraction than SCC, reflected by a smaller Outside compartment. Within BCC, NOD BCC niches were primarily composed of mCAF-associated cells, whereas INF BCC showed a shift toward myoCAF dominance, with mCAF as the next largest component. In contrast, SCC tumors (WD and PD) displayed a majority Outside fraction and a comparatively even distribution across CAF niche types among niche-associated cells; WD SCC showed a modest increase in iCAF representation relative to PD SCC (**Fig. 5D,F**).

Across cell types (**Fig. 5E**), epithelial cells were predominantly classified as Outside, but the minority of niche-associated epithelial cells mapped mainly to mCAF and myoCAF niches. Endothelial cells showed the strongest CAF neighborhood association (lowest Outside fraction), with niche-assigned endothelial cells distributed primarily across mCAF and myoCAF niches and a smaller retFIB component; iCAF and mixed contributions were minimal. Among immune lineages, B cells and neutrophils were most frequently Outside, whereas CD4 T cells, CD8 T cells, Tregs, macrophages, and NKT cells showed higher niche association and preferentially mapped to mCAF niches with variable additional contributions from myoCAF and retFIB. Mixed niches remained consistently rare, and mCAF/myoCAF accounted for the dominant share of niche-assigned cells across lineages (**Fig. 5E**).

We next asked whether these CAF-defined microenvironments were associated with distinct functional states of neighboring immune and epithelial cells. To this end, we quantified lineage-restricted marker intensities across CAF niches and the Outside compartment (**Fig. 5G**). iCAF niches emerged as inflammatory, checkpoint-high immune microenvironments. Across CD4 T cells, CD8 T cells, and Tregs, lymphocytes located in iCAF niches showed increased activation/proliferation-associated marker expression together with elevated checkpoint/exhaustion-associated markers, consistent with an antigen-experienced but potentially functionally restrained immune state. In contrast, lymphocytes localized to myoCAF niches showed a coordinated reduction in activation-, checkpoint-, and exhaustion-associated programs, indicating an immune-repressed local state. This was particularly evident in the CD8 T cell compartment, where effector-associated features, including Granzyme B, were attenuated in myoCAF-associated regions relative to iCAF and mCAF niches (**Fig. 5G**).

Macrophage programs also varied with CAF niche identity. Macrophages within iCAF niches showed a relative shift toward a CD163-high immunoregulatory/M2-like phenotype compared with myoCAF niches, exemplified by highest CD163 levels in iCAF and lowest CD163 in myoCAF, with Outside macrophages intermediate. In parallel, PD-L1 expression was highest Outside, intermediate in iCAF, and lowest in myoCAF, suggesting that macrophage checkpoint-associated programs are not uniformly CAF-associated but are particularly reduced in myoCAF-rich microenvironments (**Fig. 5G**).

Epithelial phenotypes further supported the spatial specialization of CAF niches. Epithelial Ki67 was higher in mCAF and myoCAF niches than in iCAF, retFIB, and Outside regions, indicating that mCAF– and myoCAF-associated microenvironments preferentially captured proliferative tumor epithelial cells. This pattern is consistent with the localization of these CAF niches near tumor-stroma interfaces, where proliferative keratinocyte carcinoma cells are frequently enriched. Thus, whereas iCAF niches were most strongly associated with inflammatory immune activation, mCAF and myoCAF niches were more closely linked to proliferative epithelial compartments and tumor-border-associated stromal organization (**Fig. 5G**).

Finally, direct pairwise comparisons between CAF niche classes showed that iCAF and mCAF niches differed only modestly in their associated immune-marker programs, whereas myoCAF niches were characterized by broadly lower immune-marker levels across multiple immune programs (**Fig. 5H**). Consistent with the Outside-versus-niche analysis, immune activation, checkpoint, and effector-associated features were lowest in myoCAF relative to iCAF/mCAF, and macrophage polarization/checkpoint markers showed the strongest enrichment in iCAF (CD163) and Outside (PD-L1) relative to myoCAF niches (Fig. 5H). Together, these data indicate that CAF niches are not only spatially distinct but functionally polarized: iCAF niches define inflammatory, checkpoint-high immune-cell states, whereas myoCAF niches mark immune-repressed, epithelial-proximal stromal regions.

### CAF subset composition at the invasive front stratifies immune-cell activation states

To determine how CAF programs relate to immune-cell states at the tumor-stroma interface, we performed an invasive-front-aligned spatial analysis. The epithelial-stromal transition was used as an internal anatomical landmark to define the invasive front in each ROI, building on the concept that invasive tumor borders represent spatially specialized tumor–stroma ecosystems in BCC and squamous carcinomas^44,45^. After automated front detection based on epithelial density and the opposing reticular-fibroblast-enriched stromal compartment, all cells were projected onto a shared front-relative depth axis. This enabled direct comparison of cell densities and lineage-restricted marker programs across tumor-side, invasive-front, and deeper stromal/reticular-dermis regions, despite heterogeneous tumor architecture across samples (**Fig. 6A-B**). By anchoring each ROI to a common epithelial-to-reticular-dermis spatial axis, this approach allowed us to distinguish front-localized programs from differences driven simply by variable ROI orientation, tumor size, or stromal sampling.

**Figure 6.**
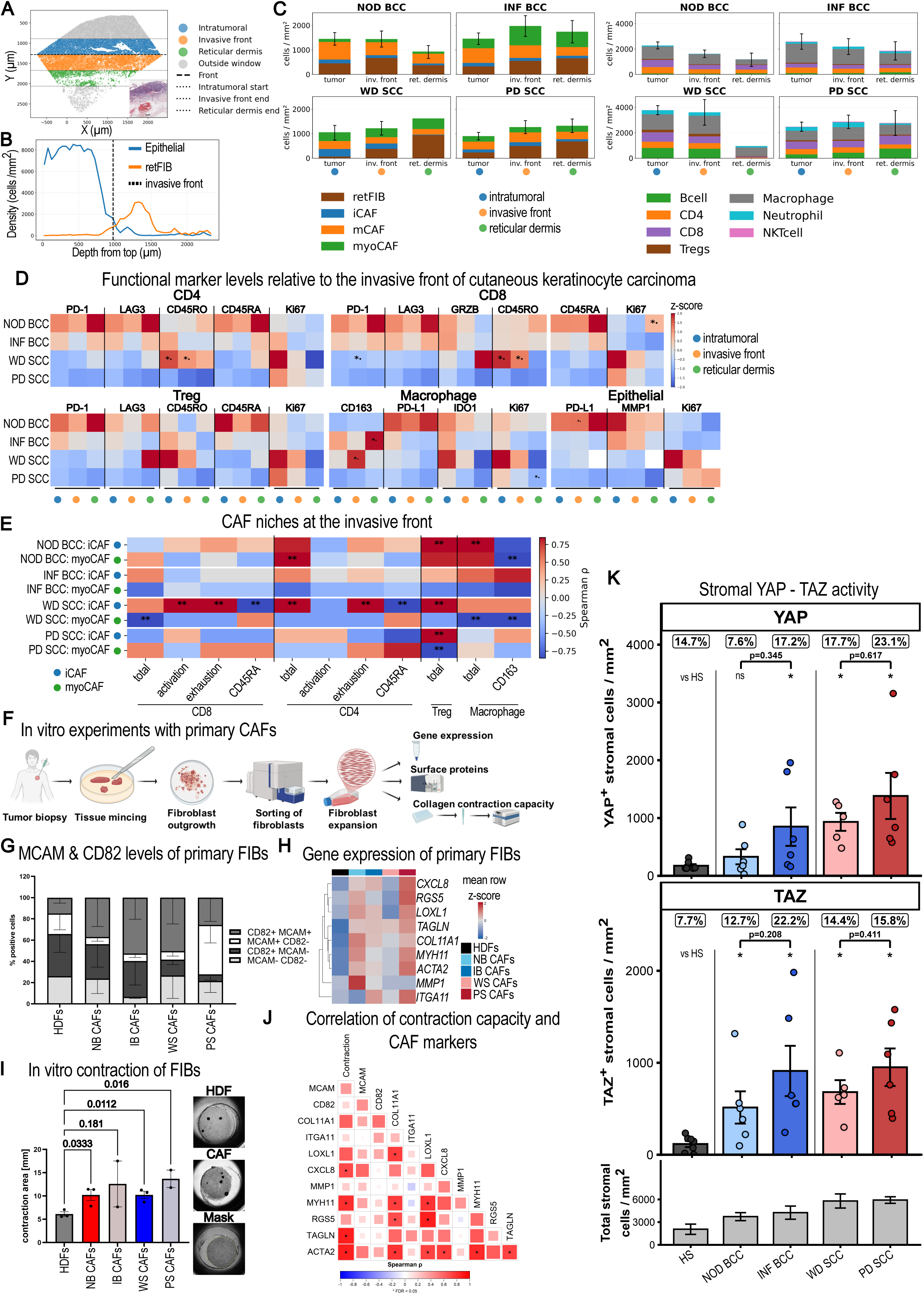
myoCAF-rich invasive fronts link immune cell exclusion to CAF contractility and YAP activation. **(A)** Example ROI after rotation into a common orientation and projection along the depth axis. Cells are assigned to three analysis compartments-intratumoral (blue), invasive front (orange), and reticular dermis (green). Dashed horizontal lines indicate the computed compartment boundaries (intratumoral start, invasive-front end, and reticular-dermis end) and the invasive-front reference line used for downstream stratification. **(B)** One-dimensional depth profiles (cells/mm²) for epithelial (blue) and reticular fibroblast (retFIB; orange) compartments in the same ROI. The vertical dashed line marks the invasive-front boundary used to separate intratumoral versus stromal depths. **(C)** Tumor-type-stratified mean cellular densities (cells/mm²; mean ± variability across samples) across compartments (intratumoral, invasive front, reticular dermis). Left: fibroblast states (retFIB, iCAF, myoCAF, mCAF). Right: immune populations (B cell, CD4, CD8, Treg, macrophage, neutrophil, NKT cell). **(D)** Marker-program heatmaps for CD4, CD8, Treg, macrophage, and epithelial cell-type sets. Each tile shows the z-scored mean arcsinh intensity (per marker, across zones) for the indicated tumor type (rows) and compartment (columns; intratumoral, invasive front, reticular dermis). Asterisks indicate within-family differential expression tested by two-sided Mann–Whitney U at the sample level (NOD BCC vs INF BCC; WD SCC vs PD SCC) and are placed only on the tumor type with the higher mean for that marker-zone combination (trend symbols, when shown, indicate 0.05-0.10). **(E)** Heatmap shows Spearman correlations (ρ) between invasive-front densities of iCAF and myoCAF (rows; two rows per tumor entity) and immune endpoints (columns) computed at the sample level within each tumor entity (NOD BCC, INF BCC, WD SCC, PD SCC). Immune endpoints (fixed order) include CD8 density, CD8 activation, CD8 exhaustion, CD8 CD45RA; CD4 density, CD4 activation, CD4 exhaustion, CD4 CD45RA; Treg density; macrophage density; macrophage CD163. Cell densities were computed as log1p(cells/mm²) within the PERI zone (normalized depth 0–0.2). Immune state scores were derived from arcsinh-transformed marker intensities (cofactor 5), aggregated to sample means within PERI, z-scored within tumor entity, and combined into composites: activation = z(CD45RO)+z(Ki67) (plus z(GranzymeB) for CD8 when available) and exhaustion = z(PD-1)+z(LAG3); CD45RA and macrophage CD163 are shown as individual z-scored markers. Asterisks denote FDR-adjusted significance within each tumor entity across all tested CAF×endpoint pairs (* q<0.10; ** q<0.05). Positive ρ indicates co-enrichment/co-activation of CAF and immune features at the invasive front; negative ρ indicates opposing spatial programs. **(F)** Graphical overview of the experimental workflow used to isolate, expand, and functionally characterize primary CAFs from patient tumor samples. **(G)** Percentages of MCAM⁺ and CD82⁺ cells in in vitro-expanded CAF cultures measured by flow cytometry. MCAM and CD82 mark myoCAF-like and iCAF-like states, respectively. **(H)** Heatmap showing relative RNA expression of selected CAF-associated genes in primary healthy dermal fibroblasts (HDFs) and patient-derived CAF cultures from the indicated tumor subtypes, determined by qPCR. **(I)** Collagen-gel contraction assay using in vitro-expanded HDFs and patient-derived CAF cultures from the indicated tumor subtypes. Representative gel images are shown on the right. **(J)** Spearman correlation matrix integrating collagen-gel contraction capacity, flow-cytometry-derived percentages of MCAM⁺ and CD82⁺ cells, and qPCR-based RNA expression of selected CAF-associated genes in the investigated primary fibroblast cultures. Asterisks indicate significant correlations after Benjamini-Hochberg correction; *FDR < 0.05. **(K)** Stromal YAP/TAZ activity in healthy skin and keratinocyte carcinoma subtypes. Quantification of nuclear YAP^+^ and TAZ^+^ stromal cells in healthy skin (HS), nodular BCC (NOD BCC), infiltrative BCC (INF BCC), well-differentiated SCC (WD SCC), and poorly differentiated SCC (PD SCC). Colored bars show the mean density of YAP^+^ or TAZ+ stromal cells per mm^2^, error bars indicate SEM, and points represent individual samples. Numbers above the colored bars indicate the mean percentage of stromal cells positive for nuclear YAP or TAZ in each group. Grey bars below show the mean total stromal cell density per mm^2^. Statistical testing was performed using planned two-sided Wilcoxon rank-sum tests comparing HS with each tumor subtype, NOD BCC with INF BCC, and WD SCC with PD SCC. P values were adjusted within each readout using the Benjamini-Hochberg method. Significance labels and p values shown in the plot refer to YAP^+^ or TAZ^+^ stromal cell density per mm^2^.

Zone-wise density profiling confirmed that the invasive front represents a distinct epithelial-stromal transition compartment. Epithelial density declined across this boundary, whereas fibroblast density increased, indicating that the front is a stromal-rich interface where tumor cells, CAFs, and immune cells converge (**Fig. 6B-C**). CAF subsets also showed compartment-specific organization in BCCs, with mCAFs appearing most prominent intratumorally in NOD BCC and myoCAFs preferentially accumulating at the invasive front of INF BCC. We therefore interpreted front-aligned marker patterns in relation to local cell availability, because immune activation, retention, or exclusion at this interface may shape the functional organization of the tumor border.

Across lymphoid compartments, front-aligned heatmaps revealed tumor-subtype-specific spatial polarization of immune-cell states (**Fig. 6D**). In WD SCC, CD4 T cells showed a tumor-proximal activated/proliferative phenotype, with similar patterns observed in CD8 T cells and Tregs. CD45RO and Ki-67 were enriched toward intratumoral and tumor-facing regions, consistent with a memory-skewed, cycling T-cell program concentrated near the tumor interface. In contrast, NOD BCC exhibited a deeper stromal shift in inhibitory and differentiation-associated markers, with PD-1, LAG3, and CD45RA enriched in reticular-dermis regions rather than at the immediate tumor-facing front. These findings are consistent with prior spatial studies showing that tumor-core and leading-edge regions can differ in transcriptional state, cellular composition, and cell–cell interactions^45^.

Myeloid and epithelial marker programs further supported subtype-specific organization of the invasive front. Macrophage CD163 was increased in INF BCC compared with NOD BCC and was enriched at the invasive front in WD SCC compared with PD SCC, consistent with localized accumulation of CD163⁺ immunoregulatory macrophage states. In contrast, macrophage PD-L1 was highest in NOD BCC, indicating a distinct checkpoint-associated macrophage program in this subtype. Tumor-cell-associated signals also differed across entities: epithelial PD-L1 was highest in NOD BCC, epithelial MMP1 was higher in BCC than SCC, and epithelial Ki-67 was higher in SCC than BCC, consistent with stronger proliferative activity in SCC and a more remodeling-associated epithelial program in BCC (**Fig. 6D**).

Because the invasive front is also a fibroblast-rich compartment, we next asked whether front-localized CAF subset composition was linked to immune-cell abundance and functional state. We therefore performed sample-level correlations between invasive-front iCAF and myoCAF densities and front-associated immune-cell densities and marker programs (Fig. 6E). By focusing on the two CAF subsets with the most divergent immune associations, this analysis tested whether immune organization at the invasive front was linked to CAF subtype composition rather than fibroblast enrichment alone.

CAF-immune coupling at the invasive front was strongly subset-dependent (**Fig. 6E**). iCAF density was associated with higher CD8 T cell activation and checkpoint/exhaustion-associated markers, increased CD4 T cell checkpoint/exhaustion-associated markers, and higher CD4 T-cell and Treg densities. In parallel, iCAF density was inversely associated with CD8 and CD4 CD45RA, consistent with enrichment of antigen-experienced, cycling/checkpoint-marked T-cell states rather than a CD45RA-high differentiation state. This pattern is consistent with the immunomodulatory role of iCAF programs previously described in skin cancer^18^. By contrast, high myoCAF density showed the opposite pattern, with negative associations with CD8 T-cell density and macrophage density, supporting a myoCAF-linked immune-low invasive-front niche. This is in accordance with studies in other epithelial cancers showing that invasive-front CAFs, particularly αSMA⁺ fibroblast-rich regions, can associate with reduced CD4/CD8 T-cell infiltration, reduced granzyme B⁺ immune cells, and CD163⁺ macrophage programs^46^.

Entity-stratified analyses showed that these CAF-immune relationships were context-dependent but remained structured. In NOD BCC, CAF enrichment was positively associated with CD4 T-cell, Treg, and macrophage densities, while inversely associated with macrophage CD163, indicating that CAF-rich front regions can coincide with immune accumulation without a parallel increase in CD163-high macrophage polarization. In INF BCC, however, myoCAF density was linked to a pronounced immune-negative axis, with broadly negative correlations across lymphoid and myeloid densities, whereas iCAF associations were comparatively weaker. PD SCC also showed divergence between CAF programs, with iCAF density aligning with Treg density and myoCAF density showing an opposing trend and stronger association with the CD45RA axis. Together, these patterns indicate that invasive-front immune organization depends on the composition of CAF subsets present at the interface, rather than on fibroblast enrichment per se.

Overall, these data support a model in which the invasive front is not a uniform boundary but a CAF-structured stromal interface. iCAF-rich front regions are linked to antigen-experienced and checkpoint-marked immune states, whereas myoCAF-rich front regions are associated with reduced immune abundance and a more immune-repressed local context. Thus, automated alignment to an epithelial-to-reticular-dermis depth axis reveals that CAF subset composition at the invasive front stratifies local immune activation states and may contribute to subtype-specific differences in immune containment versus immune escape.

### myoCAFs are contractile CAFs enriched in aggressive skin cancer subtypes

To determine whether the myoCAF-rich stromal programs identified in tumor tissues correspond to functional properties of patient-derived fibroblasts, we established an ex vivo workflow linking tumor sampling, fibroblast isolation, in vitro expansion, phenotypic profiling, transcriptional characterization, and collagen-gel contraction assays (Fig. 6F). Fibroblasts were isolated from HS (n=3) and tumor biopsies (n=10; 3× NOD BCC, 2× INF BCC, 3× WD SCC, 2× PD SCC). CAF outgrowth cultures were purified by fluorescence-activated cell sorting (FACS) using epithelial markers (EpCAM, ITGA6/CD49f, E-Cadherin) in combination with fibroblast markers (FAP, CD90). With this panel, two clearly separable populations were consistently detected, enabling exclusion of epithelial contaminants and enrichment of fibroblasts. Following FACS-sorting and expansion, fibroblast cultures were immunophenotyped by flow cytometry for CD82 and MCAM (**Fig. 6G**), marking iCAF– and myoCAF-like states, respectively, and were further characterized by quantitative PCR (qPCR) (**Fig. 6H, Supplementary Fig. 4H**). In addition, PDGFRα and Podoplanin (PDPN) protein levels were assessed by flow cytometry. Notably, all *in vitro*-expanded passage 3-4 CAF cultures were uniformly positive for both markers (data not shown), consistent with previous reports showing that the expression of various fibroblast markers changes in vitro, while the functional properties of fibroblast subsets are retained^41^. CD82/MCAM co-staining revealed marked inter-sample differences in CAF subtype composition. Healthy dermal fibroblasts (HDFs) were predominantly CD82⁺/MCAM⁻, whereas tumor-derived CAF cultures contained variable fractions of MCAM⁺ cells, including a prominent MCAM⁺/CD82⁻ subset consistent with a myofibroblast-like phenotype. CAFs derived from clinically more aggressive entities were enriched for MCAM⁺ cells, whereas CAFs from less aggressive entities retained larger CD82⁺/MCAM⁻ fractions (**Fig. 6G**). Transcript-level profiling supported these phenotypic differences. Relative to HDFs, CAF cultures showed increased expression of *CXCL8*, *COL11A1*, and contractility-associated/myoCAF-related genes (*TAGLN*, *MYH11*, *ACTA2*), while *RGS5*, *ITGA11*, and *MMP1* displayed more heterogeneous expression across samples (**Fig. 6H**, **Supplementary Fig. 4H**). Functional phenotyping using collagen gel contraction assays confirmed a tumor-associated increase in contractile capacity (**Fig. 6I**). Compared with HDFs, tumor-derived CAFs contracted collagen gels more strongly overall, with the most pronounced contraction observed in CAFs isolated from INF BCC and PD SCC (**Fig. 6I**). To integrate functional output with subtype composition and transcriptional programs, we performed correlation analyses (**Fig. 6J**). Contraction capacity correlated most strongly with contractility-associated genes, including *ACTA2*, *TAGLN*, and *MYH11*, and showed a positive trend with the fraction of MCAM⁺ cells, whereas the fraction of CD82⁺ cells, *ITGA11*, and *MMP1* were not associated with contraction. Consistently, fibroblast cultures stratified by a median split of the MCAM⁺ fraction showed significantly higher contraction capacity in the MCAM-high group, supporting MCAM as a marker of myoCAF-enriched, contractile CAF cultures (**Supplementary Fig. 4I**). Notably, RGS5 did not cluster with other myoCAF-associated markers and was detected only at very low levels in expanded fibroblast cultures, with Ct values >35 and ΔCt values of approximately 14 relative to GAPDH. This low transcript abundance mirrored our RNA-FISH observations in tumor sections, where RGS5 showed only weak in situ signal, indicating limited utility of RGS5 for identifying myoCAFs in tissue. Collectively, these ex-vivo data validate MCAM and CD82 as practical markers to resolve CAF subtype composition and link MCAM enrichment to increased contractile function.

### Stromal nuclear YAP/TAZ enrichment marks aggressive BCC and SCC subtypes and aligns with mechanotransduction-associated fibroblast programs

Mechanosensing and tissue mechanical load are closely linked to Hippo pathway signaling, with YAP and TAZ acting as key transcriptional effectors. To relate the contractile CAF program to in situ mechanotransduction, we quantified nuclear YAP and TAZ in stromal regions by multiplex immunofluorescence in an independent patient cohort comprising HS (n = 7), NOD BCC (n = 6), INF BCC (n = 6), WD SCC (n = 5), and PD SCC (n = 6) samples (**Fig. 6K, Supplementary Fig. 5A**). This cohort was analyzed separately from the IMC dataset and was used as an orthogonal validation of stromal YAP/TAZ-associated activity. K14 staining was used to delineate epithelial/tumor regions, and quantification was restricted to the surrounding stromal compartment.

Stromal YAP⁺ and TAZ⁺ cell densities were increased in tumor-associated stroma compared with HS, with the strongest elevations observed in aggressive tumor subtypes. INF BCC and PD SCC showed significant increases compared with HS for both YAP⁺ and TAZ⁺ stromal cell density, while the within-entity comparisons NOD BCC versus INF BCC and WD SCC versus PD SCC showed the same directionality but did not reach statistical significance under the current sample size (**Fig. 6K**). Because YAP⁺ and TAZ⁺ stromal cells per mm² can be influenced by the total number of stromal cells present in the tissue, we also quantified the percentage of stromal cells positive for nuclear YAP or TAZ and displayed total stromal cell density in the same panel. This allowed us to distinguish increased absolute numbers of YAP/TAZ-positive stromal cells from an increased fraction of stromal cells with nuclear YAP/TAZ positivity.

To further control for stromal cellularity, we calculated density-adjusted residuals from models regressing logit-transformed YAP⁺ or TAZ⁺ stromal-cell fractions on total stromal cell density (**Supplementary Fig. 5B**). After this adjustment, INF BCC retained higher YAP and TAZ stromal positivity than NOD BCC, whereas the SCC comparison was weaker and more variable. In tumor-only linear models including tumor entity, aggressive subtype status, and total stromal cell density, aggressive subtype status showed positive coefficients across YAP and TAZ readouts, most prominently for YAP. In contrast, total stromal cell density was most strongly associated with the cells/mm² density readout, particularly for YAP⁺ stromal cells. Together, these analyses indicate that increased stromal YAP/TAZ positivity in aggressive tumor contexts is partly linked to increased stromal cellularity but is not explained by cellularity alone.

To relate these tissue-level YAP/TAZ patterns to fibroblast-state-specific signaling programs, we reanalyzed the fibroblast populations from our published GSE254918 skin cancer single-cell RNA-sequencing dataset. PROGENy pathway inference showed that iCAFs were characterized by inflammatory signaling programs, including JAK-STAT and TNFα-related activity, whereas mCAFs showed stronger TGFβ-associated activity, consistent with their matrix-associated phenotype (**Supplementary Fig. 5C**). In the same dataset, Hippo/YAP-associated WikiPathway gene-set scoring showed the strongest mechanoregulation-associated score in RGS5⁺ cells, the mixed pericyte/myoCAF-like population described in the original study (**Supplementary Fig. 5D**). Because these WikiPathway gene sets contain a mixture of upstream regulators, pathway components and downstream response genes, we next scored selected YAP/TAZ-associated and mechanoregulatory modules that separated published YAP/TAZ response signatures and a curated YAP/TAZ–TEAD core module from curated ECM/CAF matrix and mechanotransduction-input programs (**Supplementary Fig. 5E**). This analysis showed increased ECM/CAF matrix scores in mCAFs, whereas mechanotransduction-input scores were highest in RGS5⁺ myofibroblast-like cells. In contrast, published YAP/TAZ response signature and the curated YAP/TAZ–TEAD core module were not uniformly increased across CAF populations compared with papillary fibroblasts (papFIB). Thus, ECM-remodeling and mechanotransduction-input programs were not simply mirrored by a CAF-wide increase in classical YAP/TAZ transcriptional signatures, suggesting that these modules capture distinct but related layers of stromal mechanoregulation (**Supplementary Fig. 5E**). Together with the multiplex IF data, these findings support a mechanically remodeled stromal environment in aggressive tumors, in which increased stromal nuclear YAP/TAZ coincides with complementary stromal programs, including mCAF-associated matrix remodeling and RGS5⁺/myoCAF-like mechanotransduction.

## Discussion

In this study, we used high-plex imaging mass cytometry to generate a spatially resolved atlas of basal cell carcinoma, cutaneous squamous cell carcinoma, and their surrounding stroma. Across more than 739,000 single cells from 28 regions of interest, we show that CAF heterogeneity in cutaneous carcinomas is not only compositional, but spatially organized into distinct immune, epithelial, and stromal microenvironments^8,10,13,15^. The main conceptual advance is that CAF subset identity, location within the tumor tissue, and local multicellular context are tightly linked: iCAFs identify inflamed immune-rich niches, mCAFs organize matrix-remodeling tumor-border regions where immune cells accumulate at the stromal interface, and myoCAFs define contractile, fibroblast-dense, immune-poor microenvironments associated with aggressive tumor architecture.

One of the clearest biological patterns in our data is the stromal shift that accompanies infiltrative/sclerosing BCC. Nodular BCCs were enriched for mCAF-associated stromal organization, whereas the infiltrative/sclerosing BCCs included in this cohort showed increased myoCAF representation together with higher stromal area and greater extracellular matrix deposition. This supports a model in which conventional infiltrative/sclerosing BCC is accompanied by a shift from a nest-associated matrix-producing fibroblast program toward a more contractile and desmoplastic stromal state^3,8,47,48^. Importantly, our infiltrative BCC group did not include micronodular BCCs. This distinction is relevant because micronodular BCC may be grouped together with infiltrative and sclerosing BCC in some diagnostic classification schemes^37^. Therefore, our findings should be interpreted as evidence for a myoCAF-rich, contractile stromal program in infiltrative/sclerosing BCC rather than as a general feature of all histologically high-risk BCC growth patterns. By contrast, SCC showed less dramatic subtype-specific differences in CAF composition, even though poorly differentiated SCC shared architectural features of aggressive disease, including stromal expansion and increased ECM deposition. Taken together, these findings suggest that stromal evolution is entity-specific: BCC progression appears especially tightly coupled to qualitative remodeling of CAF state, whereas SCC progression is associated with broader changes in stromal, vascular, and immune cell abundance.

A second key point is that protein-level analysis refines CAF classification in ways that transcript-based definitions alone may miss. In our dataset, myoCAFs were best resolved by αSMA and TAGLN protein expression, whereas ACTA2 RNA did not closely track with αSMA protein in situ. This is important because many single-cell studies define myoCAFs operationally by ACTA2 transcripts^8,10,13,15^, which may collapse contractile COL11A1-positive states into broader fibroblast classes. Our data therefore suggest that contractile fibroblast identity in skin cancer is not fully captured by RNA alone and that protein-level markers are required for robust distinction of matrix-producing from truly myofibroblastic CAF programs in situ^8,48^. This distinction is both methodologically and biologically relevant, as it may explain the overlap between IMC-defined myoCAFs and RNA-FISH-defined mCAFs.

Trajectory analysis further indicates that iCAFs arise via an mCAF intermediate, consistent with Forsthuber et al., 2024, and supports mCAFs as a precursor state for iCAFs^10,13,18^. Transcriptionally, iCAFs retain an mCAF-like gene expression program but show reduced expression of mCAF-associated genes together with induction of inflammatory genes absent in mCAFs. For the myoCAF lineage, the trajectory analysis suggests that an mCAF subpopulation with increased similarity to myoCAFs may represent a transitional precursor state. Compared with mCAFs, myoCAFs diverge more strongly at the transcriptomic level, showing enrichment of mural/perivascular programs and lower PDGFRα expression^8,18,24^. In situ, however, they are most reliably identified at the protein level, where myofibroblast-like programs are elevated despite persistent expression of the mCAF marker COL11A1.

The spatial analyses further show that CAF subsets occupy distinct immune microenvironments. iCAFs positively correlated with multiple immune lineages and localized to lymphocyte-rich neighborhoods, consistent with inflamed immune-rich stromal regions with elevated activation– and checkpoint-associated marker programs. mCAFs showed a different spatial pattern, associating with immune accumulation in the stromal compartment, particularly macrophages, while showing weaker or inverse relationships with immune-cell densities inside tumor nests. This supports a role for mCAF-rich regions in stromal immune compartmentalization at the tumor–stroma interface. In contrast, myoCAFs were broadly underrepresented in immune-cell-rich regions and instead enriched in fibroblast-dense, CAF-centered niches. These patterns are consistent with studies linking TGFβ-associated, matrix-rich, or myofibroblast-like stromal programs to T cell exclusion, fibroblast-dependent T cell compartmentalization, and reduced immunotherapy responsiveness^14,49–52^. Together, these findings position myoCAFs as a structurally and immunologically distinct stromal state, enriched in fibroblast-dense niches and spatially separated from immune-cell-rich regions. Thus, fibroblast abundance alone does not define tumor immune contexture; rather, CAF identity and spatial positioning appear to be key determinants of local immune organization.

Neighborhood-enrichment analysis showed that histological subtype differences were reflected in the spatial arrangement of cell types, rather than only in their abundance. NOD BCC and WD SCC retained stronger epithelial-associated neighborhoods, including epithelial-endothelial and epithelial-fibroblast associations. In contrast, INF BCC and PD SCC showed stronger immune-myeloid and selected immune-CAF neighborhood patterns, consistent with a shift toward stromal immune cell compartmentalization in more aggressive histological subtypes. Importantly, these spatial differences did not simply mirror cell-type abundance, as several subtype-associated neighborhood enrichments were not accompanied by proportional changes in cell density. Thus, CAF-associated tumor aggressiveness appears to involve remodeling of local tissue architecture, particularly at epithelial-stromal and immune-CAF interfaces, rather than only expansion or depletion of specific cell populations.

The niche analysis adds an important functional layer to this organization. T cells localized within iCAF niches displayed increased activation, proliferation, and checkpoint/exhaustion-associated marker expression, indicating that iCAF-rich regions represent inflamed but potentially restrained immune microenvironments rather than uniformly effective anti-tumor niches (Fig. 5G-H, Supplementary Fig. 4G). By contrast, myoCAF niches were associated with broadly reduced lymphocyte activation and cytotoxic features, together with lower macrophage checkpoint– and polarization-associated marker levels compared with iCAF-associated regions. These findings support a spectrum of CAF-linked immune states: iCAFs coexist with antigen-experienced, checkpoint-high immunity; mCAFs align with tumor-border and stromal immune compartmentalization; and myoCAFs define immune-repressed, fibroblast-dense microenvironments. This interpretation is consistent with studies linking matrix-rich or myofibroblast-associated stromal programs to T-cell exclusion and reduced immunotherapy responsiveness^14,49–52^. A related CAF-subset pattern was observed at the invasive front, where iCAF density was associated with activated and antigen-experienced T-cell states, whereas myoCAF density was associated with reduced immune-cell abundance. Together, these data suggest that CAF-defined stromal programs may shape whether the tumor-stroma interface adopts an immune-engaged or immune-repressed organization.

The in vitro experiments further support the biological interpretation of the myoCAF program. Primary CAFs isolated from tumors were profiled at the RNA and protein levels and functionally assessed in collagen-gel contraction assays, showing that myoCAF-enriched CAF outgrowths are more contractile, consistent with prior reports that CAFs can acquire highly contractile phenotypes and remodel the extracellular matrix^38,53^. This functional readout links the tissue-defined myoCAF program to a measurable mechanical phenotype, supporting the interpretation that myoCAF-rich tumors are not defined by marker expression alone.

In parallel, stromal nuclear YAP/TAZ was increased in aggressive tumor contexts in an independent patient cohort, supporting a mechanoregulated stromal phenotype^39^. Because YAP⁺ and TAZ⁺ stromal cell densities can be influenced by total stromal cellularity, the combined analysis of positive-cell density, percentage-positive stromal cells, and density-adjusted residuals was important to distinguish increased stromal abundance from YAP/TAZ positivity beyond cellularity alone. These analyses indicate that stromal cellularity contributes to the area-normalized YAP/TAZ signal but does not fully explain it, particularly in INF BCC.

The single-cell pathway analyses refined this interpretation by separating inflammatory, matrix-associated, and mechanotransduction-associated fibroblast programs. PROGENy pathway inference confirmed the expected inflammatory signaling profile of iCAFs and TGFβ-associated activity in mCAFs, whereas Hippo/YAP-associated gene-set analysis highlighted a mechanoregulation-associated program in RGS5⁺ cells, the mixed pericyte/myoCAF-like population from our previously published skin cancer fibroblast atlas. To further separate upstream mechanoregulatory programs from downstream YAP/TAZ transcriptional output, curated module scoring showed that mCAFs were enriched for an ECM/CAF matrix program, whereas RGS5⁺ cells showed the strongest mechanotransduction-input scores. By contrast, canonical YAP/TAZ–TEAD target-gene scores were not uniformly increased in CAF populations. This distinction is important, because nuclear YAP/TAZ protein positivity in tissue and YAP/TAZ–TEAD target-gene expression in scRNA-seq represent related but non-identical readouts. This is consistent with recent dermal fibroblast data showing that fibroblast Yap/Taz regulates collagen production, matrisome composition and higher-order ECM organization, while collagen regulation may occur indirectly through CCN2 and TGFβ/Smad rather than through direct activation of collagen genes^54^. Together, the in vitro contraction assay, stromal YAP/TAZ multiplex IF analysis, and fibroblast-subset pathway analysis support a mechanically remodeled stromal program in aggressive tumors, distributed across complementary stromal states: mCAFs as ECM/matrix-remodeling cells and RGS5⁺/myoCAF-like cells as contractile, mechanotransduction-associated cells.

The translational implication is that not all fibroblast-rich tumors are equivalent. iCAF-enriched and checkpoint-high niches may represent inflamed but restrained immune microenvironments, whereas myoCAF-dominant tumors may represent mechanically remodeled, immune-repressed contexts in which stromal reprogramming strategies could help restore immune access. In this framework, CAF subset composition is not just a descriptive histologic feature but becomes a candidate marker of immune topology and potential therapeutic vulnerability^49,50,52^. Although our data do not yet test treatment response directly, they provide a mechanistic rationale for stratifying keratinocyte carcinomas by CAF subsets, spatial organization, and activation states rather than by tumor subtype alone.

This study also has clear limitations. The cohort is modest and cross-sectional, and the analyses were not powered around predefined effect sizes. Because this study was designed as an exploratory spatial discovery study, statistical comparisons were interpreted as support for biologically coherent patterns rather than as confirmatory hypothesis tests. ROI-based sampling may enrich for morphologically informative tumor-stroma interfaces but may not capture the full heterogeneity of entire lesions. In addition, neighborhood analyses based on centroid proximity infer local spatial organization but cannot establish direct cell-cell communication or causality. The in vitro fibroblast assays support the contractile nature of myoCAF-enriched cultures, but culture conditions may reshape fibroblast states. The YAP/TAZ validation cohort was independent from the IMC cohort, which strengthens orthogonal validation but prevents direct sample-level modeling of IMC-defined myoCAF abundance against stromal YAP/TAZ activity. Finally, although orthogonal validation with RNA-FISH, IHC, multiplex immunofluorescence, scRNA-seq pathway analysis, and functional assays strengthens the conclusions, outcome-linked and longitudinal datasets will be needed to determine whether CAF-defined niches predict recurrence, metastasis, or response to immunotherapy.

Overall, this study supports a model in which cutaneous carcinomas contain spatially distinct CAF niches with divergent immune and mechanical features. In this model, iCAFs mark inflamed, immune-rich niches with elevated activation– and checkpoint-associated marker programs; mCAFs are linked to tumor-border organization and stromal immune compartmentalization; and myoCAFs define contractile, mechanosensitive, immune-poor stromal microenvironments associated with aggressive tumor architecture. By integrating IMC, RNA-FISH, IHC, multiplex IF, scRNA-seq pathway analysis, and functional contraction assays, this study identifies myoCAF-rich stromal niches as mechanically remodeled and immune-poor tumor microenvironments.

## Methods

### Ethical approval

The present study complies with all relevant ethical regulations at the Medical University of Vienna and was approved by the Institutional Review Board under the ethical permits EK#1695/2021 and EK#1783/2020. Written informed patient consent was obtained before tissue collection in accordance with the Declaration of Helsinki. Consent to publish clinical information potentially identifying individuals was obtained and approved by the data-clearing committee of the Medical University of Vienna.

### Tissue samples and ROI selection

Seventeen tumor sections were analyzed, including nine BCC samples (four nodular and five infiltrative/sclerosing) and eight SCC samples (three well differentiated and five poorly differentiated). H&E-stained sections from the same tissue blocks were used to review histopathology and representative tissue morphology, and all diagnoses and subtype assignments were confirmed by a dermatopathologist. Micronodular BCCs were not included in the infiltrative/sclerosing BCC group analyzed in this study. A total of 28 ROIs were acquired from 17 tumor samples, covering approximately 136 mm² of tissue, including 50 mm² tumor and 86 mm² stroma. ROIs were selected to capture representative tumor nests, stromal regions, and tumor-stroma interfaces.

### Antibody panel and staining

A 33-plex IMC panel was designed, including antibodies against pan-KRT, E-Cadherin, CD31, CD45, CD3, CD4, CD8, FOXP3, CD20, CD68, CD163, CD66b, MPO, CD15, PD-1, PD-L1, PD-L2, LAG-3, B7-H3, CD45RA, CD45RO, Granzyme B, Ki-67, Vimentin, FAP, αSMA, TAGLN, COL1, COL11A1, MMP1, IDO1, and CD56. Additionally, cell membranes were stained with ISCK1/2/3 and nuclei with iridium for cell identification and segmentation. All antibodies were titrated prior to use and validated by expected localization patterns. A complete list of materials, reagents and consumables used in this study is provided in Supplementary Table 1-2.

### Image acquisition and segmentation

IMC was performed on 4 µm FFPE sections, and images were acquired at subcellular resolution with a Hyperion Imaging System (Standard BioTools). Tissue segmentation and single-cell masks were generated using HALO (v3.6.4134.464) software with a self-trained nuclei segmentation plug (Nuclei Seq HALO AI) using the HighPlex plugin (HighPlex FL v4.2.14) and DenseNet V2 (HALO AI). Tissue compartments were stratified into three broad classes: epithelial/tumor, stroma, and vessel. The epithelial/tumor class was defined by epithelial marker expression and morphology and therefore included keratinocyte-derived tumor nests and, where present within the ROI, small areas of non-neoplastic epidermal or adnexal epithelium. These non-tumor epithelial structures were rare in the analyzed ROIs and were not evaluated as separate biological compartments. Other skin structures, including hair follicles, arrector pili muscle, and subcutaneous adipose tissue, were absent or only minimally represented in the selected ROIs and were therefore not separately annotated or included in compartment-specific analyses. Segmentation quality was confirmed visually on representative regions.

### Immunohistochemistry and CD3 quantification

Immunohistochemistry was performed on 4 µm human FFPE sections according to standard protocols. Antigen retrieval was conducted in citrate buffer, pH 6.0, and 3% BSA/PBST was used for blocking. Primary antibody against CD3 (1:200, rabbit, Abcam #ab16669, RRID:AB_443425) was diluted in 1% BSA/PBST and incubated overnight. A biotinylated goat anti-rabbit antibody (1:200, Vector BA-1000) was used as secondary antibody and incubated for 30 min at room temperature. Novocastra Streptavidin-HRP (Leica Biosystems Newcastle #RE7104) and Dako AEC+ High Sensitivity Substrate (Dako #K3469) were used for signal amplification and chromogenic development. Sections were counterstained with hematoxylin and scanned using an Aperio slide scanner.

CD3⁺ cells were quantified on scanned whole-slide images in HALO using a DenseNet V2 tissue classifier and the Multiplex IHC v3.4.9 analysis module, using a workflow adapted from our previous CAF study18. Tissue and tumor-nest regions were classified, and CD3⁺ cells were detected within the corresponding masks. CD3⁺ cell densities were calculated as positive cells/mm² for whole tissue and intratumoral/tumor-nest compartments.

### Masson’s trichrome staining

Masson’s trichrome staining (DAKO) was performed on consecutive tissue sections to visualize collagen-rich extracellular matrix and overall stromal architecture. Stained slides were scanned as whole-slide images using a Vectra Polaris multispectral imaging system. Digital images were analyzed in HALO, where representative tumor, stromal, cellular, ECM-rich, and background/non-tissue regions were manually annotated and used to train a DenseNet-based tissue classifier. The trained classifier was subsequently applied to the complete slide images. HALO area-quantification tools were used to generate tissue, cellular, and ECM masks and to quantify stromal area and ECM coverage across histological tumor subtypes.

### Multiplex immunofluorescence staining and imaging

Formalin-fixed, paraffin-embedded tissue sections from an independent patient cohort comprising healthy skin (HS; n = 7), nodular BCC (NOD BCC; n = 6), infiltrative/sclerosing BCC (INF BCC; n = 6), well-differentiated SCC (WD SCC; n = 5), and poorly differentiated SCC (PD SCC; n = 6) were used to assess stromal YAP/TAZ activity. Sections were deparaffinized, rehydrated, and subjected to antigen retrieval using standard protocols. Primary antibodies against YAP (rabbit monoclonal, Cell Signaling Technology, #14074, RRID:AB_2650491), TAZ/WWTR1 (mouse monoclonal, Sigma-Aldrich, #AMAB90730), and Keratin 14 (K14; chicken polyclonal, BioLegend, #906004, RRID:AB_2616962) were applied. Primary antibodies were detected using species-specific secondary antibodies: goat anti-rabbit IgG Alexa Fluor 594, goat anti-mouse IgG Alexa Fluor 647, and goat anti-chicken IgY Alexa Fluor 488 (Thermo Fisher Scientific). Nuclei were counterstained with DAPI.

Slides were imaged using a Vectra Polaris multispectral imaging system with identical acquisition settings across all samples. Multispectral images were analyzed in HALO. K14 staining was used to define epithelial/tumor regions, and YAP/TAZ quantification was restricted to the surrounding stromal compartment. Nuclear YAP⁺ and TAZ⁺ stromal cells were detected using a nuclear-positive cell-detection algorithm. For each sample, YAP⁺ and TAZ⁺ stromal-cell densities were calculated as positive stromal cells/mm². In addition, total stromal cell density was calculated as stromal nuclei/mm², and the percentage of stromal cells positive for nuclear YAP or TAZ was calculated as the number of YAP⁺ or TAZ⁺ stromal cells divided by the total number of stromal cells.

The YAP/TAZ analysis therefore generated three related readouts: YAP⁺ or TAZ⁺ stromal-cell density, reported as positive cells/mm²; the percentage of stromal cells positive for nuclear YAP or TAZ; and density-adjusted stromal YAP/TAZ positivity, calculated to account for differences in total stromal cellularity. Statistical testing, including group comparisons and density-adjusted residual analysis, is described in the Statistical analysis section.

### RNA-FISH/αSMA immunofluorescence co-staining

Multiplex RNA fluorescence in situ hybridization (RNA-FISH) was performed on FFPE tissue sections using the RNAscope Multiplex Fluorescent Reagent Kit v2 (ACD Bio-Techne, catalog no. 323135), including PretreatPRO pretreatment, according to the manufacturer’s instructions. For CAF subset validation on sections consecutive to those used for IMC, a 4-plex RNA-FISH panel was used to detect MMP1 (catalog no. 412641-C1), COL11A1 (catalog no. 400741-C2), RGS5 (catalog no. 533421-C3), and COL1A1 (catalog no. 401891-C4). Probes were detected by tyramide signal amplification using Opal fluorophores from Akoya Biosciences as follows: MMP1 with Opal 570 Reagent Pack (catalog no. FP1488001KT; 1:500), COL11A1 with Opal 620 Reagent Pack (catalog no. FP1495001KT; 1:500), RGS5 with Opal 690 Reagent Pack (catalog no. FP1497001KT; 1:500), and COL1A1 with Opal TSA-DIG Reagent (catalog no. OP-001007; 1:500) followed by Opal 780 Reagent (catalog no. OP-001008; 1:200), supplied as the Opal 780 Reagent Pack (catalog no. FP1501001KT). DAPI was used as nuclear counterstain. This panel was used to identify RNA-FISH-defined iCAFs and mCAFs and to visualize RGS5⁺ myoCAF-like stromal regions.

To directly compare *ACTA2* RNA with αSMA protein expression in the same tissue section, a separate 4-plex RNA-FISH assay was combined with αSMA immunofluorescence. RNA-FISH probes targeted *MMP1* (catalog no. 412641-C1), *ACTA2* (catalog no. 444771-C2), *COL11A1* (catalog no. 400741-C3), and *COL1A1* (catalog no. 401891-C4). Probes were detected using Opal fluorophores from Akoya Biosciences as follows: *MMP1* with Opal 570 Reagent Pack (catalog no. FP1488001KT; 1:500), *ACTA2* with Opal 690 Reagent Pack (catalog no. FP1497001KT; 1:500), *COL11A1* with Opal 620 Reagent Pack (catalog no. FP1495001KT; 1:500), and *COL1A1* with Opal TSA-DIG Reagent (catalog no. OP-001007; 1:500) followed by Opal 780 Reagent (catalog no. OP-001008; 1:200), supplied as the Opal 780 Reagent Pack (catalog no. FP1501001KT). After completion of RNA-FISH, sections were immunostained for αSMA using mouse anti-αSMA antibody (clone 1A4; Abcam, catalog no. ab7817; 1:2000), followed by donkey anti-mouse IgG (H+L) highly cross-adsorbed Alexa Fluor 488 secondary antibody (Thermo Fisher Scientific/Invitrogen, catalog no. A-21202; 1:300). DAPI was used as nuclear counterstain.

Images from both RNA-FISH assays were acquired using a Vectra Polaris multispectral imaging system and analyzed in HALO. RNA-FISH-stained sections were quantified using the workflow described in our previous publication. Because RGS5 expression in stromal cells was close to the detection threshold, RGS5⁺ stromal cells were used for spatial visualization but were not robustly quantified. RNA-FISH-based CAF quantification therefore focused on iCAFs, defined as COL1A1⁺MMP1⁺ cells, and mCAFs, defined as COL1A1⁺COL11A1⁺MMP1⁻ cells. αSMA immunofluorescence was used to assess protein-level contractile CAF features and to compare αSMA protein signal with ACTA2 RNA signal on the same section.

### Fibroblast isolation from tumor samples

Cancer-associated fibroblasts were isolated by outgrowth of minced tumor pieces in DMEM containing 10% FBS, Gentamicin (Gibco #15710049, 1:200) and Normocin (InvivoGen #ant-nr-05, 1:500). After 1–2 weeks depending on the growth rate, cells were harvested with Versene followed by Accutase treatment and stained for E-Cadherin/CDH1 (#324110, BioLegend, 1:100), EpCAM (#324110, BioLegend, 1:100), ITGA6/CD49f (#MCA699F, Bio-Rad, 1:20), PDGFRα/CD140a (#323508, BioLegend, 1:20), Podoplanin (#337004, BioLegend, 1:100), CD90 (#328116, BioLegend, 1:20) and FAP (#FAB3715P, R&D, 1:20). DAPI was used as live/dead marker (1:5000). Cells were sorted using a FACSAria (BD Biosciences) into epithelial cells and fibroblasts. All fibroblasts were CD90+, FAP+, PDGFRa+ and Podoplanin+. Fibroblasts were stained with MCAM/CD146 (# 550315, BD Biosciences, 1:10) and CD82 (# 564341, BD Biosciences, 1:80) to determine the percentage of immunomodulatory CAFs and myofibroblast-like CAFs. Sorted cells were expanded and frozen at passage 3. Patient information for the primary fibroblast cultures used in this study is provided in Supplementary Table 3.

### Quantitative PCR

RNA was isolated with the Qiagen RNeasy Mini Kit (Qiagen #74106). RevertAid H Minus First Strand cDNA Synthesis Kit (Thermo Scientific #K1631) was used to prepare cDNA after a DNase I digestion step (Thermo Scientific #EN0521). TaqMan 2xUniversal PCR Master Mix (Applied Biosystems #4324018) and TaqMan probes for GAPDH (Hs99999905), ITGA11 (Hs01012939), COL11A1 (Hs01097664_m1), LOXL1 (Hs00935937_m1), CXCL8 (Hs00174103_m1), MMP1 (Hs00899659_g1), RGS5 (Hs01591223_s1), ACTA2 (Hs00426835_g1), TAGLN (Hs01038777_g1), MYH11 (Hs00975796_m1) were used in the qPCR.

### Contraction assay

CAFs previously isolated from tumor samples were cultured in DMEM supplemented with 10% FBS and penicillin-streptomycin and seeded in 10 cm dishes coated with neutralized 0.1 mg/ml rat-tail collagen I (COL1; Corning, catalog no. 354236). After 24 hours, medium was exchanged to DMEM containing 2.5% FBS, TGF-β (30 pg/ml, PeproTech® #100-21-10ug), FGF (5 ng/ml, PeproTech® #100-18B), EGF (5 ng/ml, R&D Systems #234-FSE), Insulin (5 µg/ml, Sigma-Aldrich #I979278-5ML), 2-phosphate ascorbic acid (50 µg/ml, Santa Cruz Biotechnology #SC-228390), Pen-Strep (1%, Gibco #15140-122), non-essential amino acids (1X, Gibco #11140-050), Hydrocortisone Hemisuccinate (1 µg/ml, Sigma-Aldrich #H0888-1G) and glutamine (7.5 mM, Gibco #25030-024). After 6 days, 4 × 10⁴ cells were embedded in COL1-Matrigel mixture (Matrix composition: neutralized 2 mg/ml Rat-tail Collagen1 (Corning # 354236), 25% Matrigel (Corning # 356231), 1x DMEM), and seeded in 24-well plates at 200 µl per well, with at least three replicates per condition. After 1 hour the gels were punched with a 5 mm biopsy punch and the same media was added. Contraction was evaluated after 40 hours by imaging the wells with Cytation 5 at 10x magnification, and the remaining area of the contracted gels was manually measured in Fiji. Contracted area was determined by subtracting the remaining area from the area of the 5 mm biopsy punch.

### Preprocessing and clustering

Values of single-cell intensities for each marker were exported from HALO, capped at the 99th percentile, and arcsinh-transformed with a cofactor of 1 prior to clustering. Computational analyses were performed in R v4.4.0 using Seurat v5.3.0 and harmony v1.2.3. In Seurat, preprocessing included normalization, variable feature selection, data scaling, and principal component analysis. PCA was performed using 30 components, followed by batch correction with Harmony using dataset as the grouping variable. Graph-based clustering was performed using the shared nearest neighbor (SNN)/Leiden algorithm on the Harmony reduction with dimensions 1:20. Clustering was assessed at resolutions 0.1, 0.2, and 0.5, and resolution 0.5 was used for downstream analyses. UMAP was computed using dimensions 1:20 on both PCA and Harmony embeddings. Unless otherwise specified, default Seurat parameters were used.

### Fibroblast clustering and subset annotation

All non-immune, non-epithelial, non-endothelial, and CD56^-^ cells were merged and subclustered to resolve fibroblast lineages. Subclustering was performed in Seurat after normalization, variable feature selection, scaling, and PCA using 20 principal components, followed by graph-based clustering with dims = 1:20 and resolution = 0.5. Vimentin⁺ αSMA^high^TAGLN^high^ clusters were assigned as myoCAFs. The largest cluster (Vimentin⁺COL1A1⁺COL11A1⁺αSMA⁻), localized mainly in the reticular dermis and distant from tumor nests, was annotated as reticular fibroblasts. One artifactual cluster representing mis-segmented collagen fibers was excluded. The remaining clusters were further subclustered using MMP1, IDO1, FAP, and COL11A1 expression to define iCAFs (Vimentin⁺MMP1⁺ and/or IDO1⁺, αSMA^low^) and mCAFs (Vimentin⁺COL11A1⁺, MMP1⁻IDO1⁻, αSMA^low^).

### Sample-level compositional analysis

Cell-type composition was compared at the biological-sample level to avoid ROI-level pseudoreplication. ROIs from the same sample were aggregated before analysis, and planned comparisons were performed between NOD BCC and INF BCC, and between WD SCC and PD SCC, separately for all tissue, tumor, and stroma. For each sample, cell-type counts were analyzed as compositional data using centered log-ratio transformation after zero-count replacement. This approach accounts for the relative nature of cell-type abundance data, where an apparent increase in one cell type necessarily affects the proportions of others. Group differences were assessed for each cell type using two-sided Wilcoxon rank-sum tests, with Benjamini-Hochberg correction within each comparison and compartment. Effect sizes are reported as differences in median centered log-ratio abundance, with positive values indicating enrichment in INF BCC relative to NOD BCC or in PD SCC relative to WD SCC. Cell types were grouped into epithelial, immune, fibroblast, endothelial, and other major classes for visualization.

### Cell densities correlation analysis

Cell-type densities from IMC were summarized at the sample level as cells/mm² within each annotated compartment (tissue, stroma, and tumor nests). RNA-FISH CAF densities and CD3 immunohistochemistry (IHC) T-cell densities were derived from whole-section quantification and stored as cells/mm² for each sample. To stabilize variance and reduce the influence of zeros and extreme values, densities were transformed as log10(density + ε) with ε = 0.01 prior to correlation analyses. Associations between modalities were assessed using two-sided Spearman rank correlation (ρ) across samples. For IMC vs RNA-FISH CAF concordance, Spearman correlations were computed between IMC CAF subset densities and RNA-FISH CAF subset densities using both matched and alternative subtype pairings (IMC iCAF vs RNA iCAF; IMC mCAF vs RNA mCAF; IMC myoCAF vs RNA mCAF; IMC myoCAF vs RNA iCAF). For the CAF vs CD3 analysis, Spearman correlations were computed between CAF densities quantified by IMC or RNA-FISH (iCAF, mCAF, myoCAF, iCAFs.RNA, mCAFs.RNA) and CD3⁺ T-cell densities quantified by IHC in whole tissue and intra-nest compartments, both for all samples combined and stratified by tumor entity (SCC and BCC).

Within the IMC dataset, correlation analyses were performed using compartment-specific cell densities. A tissue-only lineage correlation matrix was computed from lineage densities within the tissue compartment. In addition, a compartment-aware CAF–immune correlation matrix was generated by correlating tissue CAF subset densities with immune lineage densities quantified separately in stroma and tumor nests. For these IMC heatmaps, densities were transformed using log(1 + density) prior to correlation, and two-sided Pearson correlation (r) was used.

For heatmap annotations, asterisks indicate significance thresholds based on p values; for the CAF–CD3 heatmaps in Fig. 4C, asterisks denote nominal (unadjusted) p values (*p < 0.05; **p < 0.01; ***p < 0.001). Where explicitly stated for other heatmap summaries aggregating multiple comparisons (e.g., CAF–immune blocks), p values were adjusted for multiple testing using the Benjamini-Hochberg procedure and adjusted significance is indicated accordingly. For the CAF–immune correlation matrix, Benjamini-Hochberg adjustment was applied separately within the stroma and tumor comparison families.

### Spatial analysis

Neighborhood enrichment was computed using Squidpy v1.6.1. A spatial neighbor graph was constructed from cell centroids with sq.gr.spatial_neighbors (coord_type=“generic”, radius = 100 µm), defining neighbors as pairs of cells whose centroid-to-centroid Euclidean distance was ≤100 µm. Neighborhood enrichment was then calculated with sq.gr.nhood_enrichment (cluster_key=“cell_type”), which compares observed cell–cell adjacencies to a permutation-based null model and reports a z-score per cell-type pair (observed minus permuted mean, divided by permuted standard deviation; default n_perms = 1000 in Squidpy 1.6.1).

For each ROI, the neighborhood-enrichment z-score matrix was extracted from adata.uns[“cell_type_nhood_enrichment”][“zscore”]. ROI-level matrices were averaged within each sample (mean across ROIs) to yield a per-sample neighborhood-enrichment matrix. The grand mean neighborhood-enrichment matrix across all samples (mean of per-sample matrices) was visualized as a heatmap (Fig. 4D). For each tumor subtype, mean neighborhood-enrichment matrices were generated by averaging the per-sample matrices and visualized as network graphs (Fig. 4E). Nodes represent cell types, and edges indicate cell-type pairs with neighborhood-enrichment scores of |z| ≥ 1.96. Edge color denotes enrichment (red) or depletion (blue), while edge width reflects the magnitude of the z-score. All networks were plotted using a fixed ring layout with consistent node ordering across tumor subtypes.

To relate interaction changes to compositional differences between subtypes, we calculated differential neighborhood enrichment for each comparison (NOD BCC vs INF BCC; WD SCC vs PD SCC) as the difference in subtype-mean neighborhood-enrichment z-scores for each cell-type pair. Cell-type abundance was quantified as cell density (cells/mm^2^) per ROI, derived by dividing the number of cells of each type by ROI area; ROI densities were averaged to the sample level and then averaged within subtype. Density differences were defined as mean cell density in subtype A minus mean cell density in subtype B.

For the interaction-versus-abundance plots (Fig. 4G), each point represents a cell-type pair and is positioned by cell-density difference on the x-axis and differential neighborhood enrichment on the y-axis (A−B). Points were filtered to pairs with absolute differential neighborhood enrichment above the plotting threshold; the top 25 pairs by absolute differential neighborhood enrichment were annotated. Positive values indicate stronger neighborhood association in subtype A (NOD BCC or WD SCC), whereas negative values indicate stronger association in subtype B (INF BCC or PD SCC). The accompanying bar plots (Fig. 4F) show per-cell type density differences (A-B), summarized as mean across samples.

### Spatial CAF subsets

To characterize fibroblast-centered spatial contexts, we quantified the local neighborhood composition of each marker-defined fibroblast (iCAF, mCAF, myoCAF, retFIB) within each ROI. For every fibroblast centroid, neighboring cells within a 100 µm Euclidean radius were identified using a KD-tree search, excluding the index cell itself, and neighbors were counted by major cell class (epithelial, endothelial, macrophage, neutrophil, CD4 T cell, Treg, CD8 T cell, NKT cell, B cell, and fibroblast). To define neighborhood states independently of fibroblast marker identity, all fibroblast neighbor subtypes were collapsed into a single fibroblast (FIB) category. Neighbor counts were then converted to fractions by dividing by the total number of neighbors for each fibroblast. Fibroblasts with fewer than two neighbors within 100 µm were excluded from downstream neighborhood-state analysis.

Unsupervised spatial neighborhood states were identified by applying non-negative matrix factorization (NMF) to the fibroblast-by-feature neighborhood-fraction matrix using five components (NNDSVDa initialization; max_iter = 2000). To reduce confounding by heterogeneous ROI sizes and fibroblast densities, NMF fitting incorporated row weighting that combined ROI balancing, such that each ROI contributed approximately equally, with neighbor-count weighting to downweight low-neighbor estimates. Each fibroblast was assigned to a spatial FIB state (FIB0–FIB4) according to its highest NMF loading.

Marker-defined CAF identities were mapped onto spatial CAF states by cross-tabulation, reporting both the marker composition of each spatial state and the distribution of spatial states within each marker-defined subset. Enrichment of marker-defined CAF subsets in individual spatial CAF states was assessed using one-sided Fisher’s exact tests, comparing fibroblasts of a given marker-defined subtype within a given spatial state to all other fibroblasts in a 2 × 2 contingency table. P-values were adjusted for multiple testing using the Benjamini-Hochberg false discovery rate method. Effect sizes were reported as log2 odds ratios, calculated with a Haldane–Anscombe correction (+0.5) to stabilize sparse tables, whereas Fisher’s exact p-values were left unmodified.

### RNA-protein correlation scatterplot / hexbin

Single-cell intensity data were exported from the image analysis pipeline and aggregated across all samples. Cells were restricted to CAFs (COL1A1+ cells). Marker channels were taken directly from the per-cell intensity columns Opal 520 Cell Intensity (αSMA; reported here as Opal520) and Opal 690 Cell Intensity (ACTA2; reported here as Opal690). To reduce the influence of heavy-tailed intensity distributions and zero inflation, intensities were transformed using the natural log with pseudocount (log1p), yielding log520 = log1p(Opal520) and log690 = log1p(Opal690). Correlation between channels was computed on the pooled CAF population using both Pearson’s correlation coefficient (r; linear association) and Spearman’s rank correlation coefficient (ρ; monotonic association). All analyses and plotting were performed in R using the tidyverse/ggplot2 stack. For visualization, the pooled distribution of log520 vs log690 was rendered as a hexagonal binned density plot (hexbin; 70 bins), and an ordinary least squares linear regression fit (geom_smooth with method=“lm”, no confidence band) was overlaid.

### ACTA2 & αSMA correlation analysis

Multiplex immunofluorescence (Opal) single-cell intensity measurements were exported from per-cell object results tables (HALO). CAFs and αSMA+ CAFs were defined using the existing classifier output from HALO (threshold based classification). ACTA2 RNA-FISH signal was quantified as Opal690 cell intensity, and values were log-transformed as log1p(Opal690) to reduce the influence of heavy-tailed intensity distributions. To avoid confounding by between-sample differences in staining intensity and imaging conditions, comparisons were performed using within-sample centering. Specifically, for each sample, log1p(ACTA2) values were centered by subtracting the sample-wise median across all CAFs in that file, yielding centered log1p(ACTA2 cell intensities). Samples were retained only if both αSMA+ and αSMA− CAF groups contained at least 500, excluding 2 samples from the analysis. For visualization, at most 80,000 CAFs were randomly sampled from the pooled centered dataset. For each sample, the median centered log1p(ACTA2 cell intensity) was calculated separately for αSMA+ and αSMA− CAFs. The per-file effect size was summarized as Δmedian(log1p ACTA2) = median(log1p ACTA2 | αSMA+) − median(log1p ACTA2 | αSMA−), and the reported “median Δmedian” represents the median of these per-sample Δmedian values across all included samples.

### CAF niche definition and cell assignment

CAF niches were defined using a spatial seed-disk approach based on local CAF density and subtype dominance. CAFs were identified from cell-type annotations and grouped into four subtype labels: iCAF, mCAF, myoCAF, and retFIB. Spatial positions were defined by the centroid coordinates of segmented cells. For each ROI, candidate niche seeds were evaluated for every CAF cell within a fixed radius of 50 µm. A CAF cell was considered a seed if at least 10 CAFs were present within this radius. Seed subtype was assigned according to local neighborhood composition: if the most abundant CAF subtype represented at least 50% of neighboring CAFs, the seed was assigned to that subtype; otherwise, it was labeled Mixed. Thus, Mixed reflects insufficient subtype dominance within the local CAF neighborhood and is independent of geometric overlap between niche regions.

Each seed defined a circular niche region with a radius of 50 µm centered on the seed coordinate. Because these regions could overlap, non-CAF cells were assigned to a single niche class using an exclusive assignment rule. A cell was considered inside a niche class if it fell within at least one disk of that class. If a cell fell within disks of multiple classes, it was assigned according to a fixed priority order: iCAF, mCAF, myoCAF, retFIB, and Mixed. This ensured that each non-CAF cell belonged to no more than one niche class.

Two complementary definitions of Outside were used depending on the analysis. For composition plots, cells not located within any niche disk were labeled Outside, so that the sum of all niche classes and Outside equaled 1. For marker comparisons and heatmaps, a more stringent far-Outside baseline was used. Cells were classified as far-Outside if their distance from the edge of the union of all niche disks was at least 50 µm. Cells located in the intermediate zone between niche edges and the far-Outside boundary were excluded from these comparisons to reduce boundary effects and sharpen niche-versus-Outside contrasts.

For marker analysis, marker intensities were extracted from the per-cell intensity matrix and transformed before summarization to stabilize variance. Unless otherwise specified, values were clipped at the upper 99th percentile for each marker and subsequently arcsinh-transformed using a cofactor of 1.0. ROIs belonging to the same sample were aggregated before downstream visualization and statistical analysis, so that each biological sample contributed one value per niche class, cell type, and marker. For each sample-by-niche-class combination, marker expression was summarized across cells using the median, or the mean where explicitly specified. Heatmaps were generated from these per-sample summaries.

For the CAF-column row-z heatmap, z-scores were computed for each cell type–marker combination using CAF niche columns only, including iCAF, mCAF, myoCAF, retFIB, and Mixed. The resulting scaling parameters were then applied to all columns, including Outside, to emphasize relative niche enrichment or depletion within each row. For spatial overlays, ROI coordinate systems were obtained from a per-ROI orientation table. CAF subtype points and niche disks were plotted using the corresponding spatial coordinates. Optional halo-density layers were rendered by rasterizing cell locations to a grid and applying Gaussian smoothing for visualization only.

### Invasive-front-aligned spatial analysis

For each ROI, single-cell spatial coordinates were rigidly rotated to a common orientation to standardize tissue geometry across the cohort. A continuous tissue-depth axis was then defined along the aligned vertical coordinate. When epithelial cells were present, the epidermal or tumor-facing epithelial boundary was inferred from their spatial extreme and used as the y = 0 reference; otherwise, the uppermost tissue edge was used. Depth was calculated as a non-negative distance increasing into the tissue.

Cells were binned along the depth axis and converted to continuous densities in cells/mm². To account for non-rectangular tissue geometries, bin areas were locally estimated using robust lateral tissue widths, defined by the 5th–95th percentile range of x-coordinates, and scaled such that the sum of all bin areas matched the known total physical area of the ROI. For each ROI, the invasive front was algorithmically identified from smoothed density profiles as the shallowest depth showing a coordinated decrease in epithelial density and increase in reticular fibroblast density. Manual overrides were applied for a predefined subset of architecturally exceptional ROIs. Spatial depth was then normalized by the effective ROI thickness, defined as the 2nd-98th percentile depth span, and the invasive front was set to z = 0.

Each cell was assigned to one of three front-relative zones: intratumoral/tumor-side region, defined as normalized depth −0.2 to 0; invasive-front region, defined as 0 to 0.2; and reticular-dermis/deeper stromal region, defined as 0.2 to 0.4. Cells outside this normalized window were excluded from downstream zone-specific analyses. To prevent boundary-discretization artifacts, absolute zone areas were calculated by allocating the physical area of each depth bin to zones based on the exact fraction of cells belonging to each zone within that bin. Zone-specific cell densities were then computed for each cell type as total cell count divided by the corresponding zone area, ensuring consistency between quantitative density outputs and single-cell spatial overlays. To avoid pseudoreplication from uneven ROI sampling, ROIs from the same patient/sample were pooled by summing raw cell counts and absolute zone areas before calculating final sample-level densities. Tumor-group summaries were then calculated across independent samples.

For functional marker analysis along the invasive-front axis, raw marker intensities were transformed using an inverse hyperbolic sine transformation with a cofactor of 5. ROI-level marker means were aggregated to sample-level values using cell-count weighting to prevent cell-sparse ROIs from disproportionately influencing expression estimates. Sample-level values were then averaged across independent samples within tumor groups. Heatmaps display sample-level means across marker-by-zone combinations, scaled as z-scores within each marker block to emphasize relative spatial gradients.

To quantify CAF–immune relationships at the invasive front, CAF subset abundance and immune density/state endpoints were summarized within the invasive-front zone, corresponding to normalized depth 0 to 0.2. CAF predictors were restricted to iCAF and myoCAF densities and were calculated as log1p-transformed cells/mm². Immune endpoints included log1p-transformed densities of CD4 T cells, CD8 T cells, Tregs, and macrophages, as well as sample-level immune-state scores derived from functional marker intensities. Marker intensities were arcsinh-transformed with a cofactor of 5 and aggregated to sample-level means using cell-count weighting within the invasive-front zone. Within each tumor entity, markers were z-scored across samples to place markers on a comparable scale before generating composite immune-state scores. T-cell composite states were defined as follows: CD4 activation, z(CD45RO) + z(Ki67); CD4 exhaustion, z(PD-1) + z(LAG3); CD8 activation, z(CD45RO) + z(Ki67) + z(Granzyme B), with Granzyme B omitted where unavailable; and CD8 exhaustion, z(PD-1) + z(LAG3). Differentiation state was summarized by CD45RA z-scores for CD4 and CD8 T cells. Macrophage functional state was represented by CD163 z-scores. Tregs were analyzed by density only, without additional marker-composite scores. Statistical testing of CAF–immune associations, including correlation analysis, multiple-testing correction, and significance thresholds, is described in the Statistical analysis section.

### Single-cell pathway and YAP/TAZ-associated module analysis

Single-cell RNA-sequencing data from human healthy skin, basal cell carcinoma, cutaneous squamous cell carcinoma, and melanoma samples were obtained from the previously published GSE254918 dataset and analyzed using the Seurat framework in R, as described in Forsthuber et al., 2024. Fibroblast subset annotations were retained from the original study. For the pathway and module analyses shown in Supplementary Fig. 5C–E, fibroblast/CAF populations included papFIB, retFIB, mCAF, iCAF, and RGS5⁺ cells. RGS5⁺ cells were interpreted according to the original annotation as a mixed pericyte/myoCAF-like population rather than as a pure CAF subset^18^.

For broad signaling pathway inference shown in Supplementary Fig. 5C, normalized single-cell expression data were analyzed using decoupleR with the PROGENy pathway model, which infers pathway activity from perturbation-derived pathway-responsive genes. PROGENy pathway activity scores were calculated for the available signaling pathways, summarized as mean scores across fibroblast/CAF populations, and visualized as heatmaps^55^.

For the Hippo/YAP-associated gene-set analysis shown in Supplementary Fig. 5D, gene sets were obtained from the MSigDB C2 curated collection, including WikiPathways-derived gene sets, using the msigdbr package v7.5.1. Three WikiPathways gene sets were selected: WP_HIPPOYAP_SIGNALING_PATHWAY, WP_HIPPO_SIGNALING_REGULATION_PATHWAYS, and WP_MECHANOREGULATION_AND_PATHOLOGY_OF_YAPTAZ_VIA_HIPPO_AND_NONHIPPO_MECHANISMS. Gene-set scores were calculated using decoupleR, summarized as mean scores across fibroblast/CAF populations, and visualized as heatmaps. Because these gene sets contain upstream regulators, pathway components, and downstream genes, they were interpreted as Hippo/YAP-associated gene-set scores rather than direct measurements of nuclear YAP/TAZ activity.

For the module analysis shown in Supplementary Fig. 5E, we scored one published MSigDB C6 YAP response signature together with three curated modules designed to distinguish canonical YAP/TAZ transcriptional output from upstream mechanical and matrix-associated programs. The published YAP response module corresponded to the MSigDB C6 CORDENONSI_YAP_CONSERVED_SIGNATURE^56^. The curated YAP/TAZ–TEAD core module represented canonical YAP/TAZ–TEAD target-gene output and included established YAP/TAZ-responsive genes such as CYR61/CCN1, CTGF/CCN2, ANKRD1, AMOTL2, AXL, SERPINE1, and THBS1^57^. The mechanotransduction-input module included genes associated with integrin/focal adhesion signaling, Rho/ROCK activity, actomyosin organization, and contractility^58^. The ECM/CAF matrix module included collagen, matrix-remodeling, crosslinking, and CAF-associated ECM genes^39^. Genes for the curated modules were selected a priori from published YAP/TAZ target-gene signatures, mechanotransduction studies, and CAF matrix-remodeling literature, and were not selected based on differential expression in the present dataset. Gene aliases were resolved where necessary, including CCN1/CYR61 and CCN2/CTGF, and only genes detected in the single-cell RNA-sequencing object were used for scoring.

Module scores for Supplementary Fig. 5E were calculated using AddModuleScore in Seurat. Complete input gene lists used for each module are provided in the Supplementary Table 5.

### Statistical analysis and visualization

Computational analyses were performed in R v4.4.0 and Python v3.12.4. Single-cell preprocessing and clustering were performed in R using Seurat and Harmony, whereas spatial analyses were performed in Python using Squidpy and standard scientific Python packages, including pandas and SciPy. Unless otherwise stated, statistical analyses were performed at the independent patient/sample level. For measurements derived from multiple ROIs per sample, ROIs were pooled or aggregated before statistical testing to avoid pseudoreplication.

Planned pairwise comparisons were performed within histological tumor families, including NOD BCC versus INF BCC and WD SCC versus PD SCC. Where healthy skin was included as an external reference group, healthy skin was compared with each tumor subtype. For unpaired group comparisons of cell-density measurements, marker-summary values, fibroblast culture readouts, and YAP/TAZ readouts, two-sided Mann–Whitney U tests, equivalent to Wilcoxon rank-sum tests, were used unless otherwise specified. Where multiple cell types, markers, zones, readouts, or comparisons were tested, P values were adjusted using the Benjamini-Hochberg false discovery rate (FDR) procedure within the corresponding analysis stratum.

For YAP/TAZ multiplex immunofluorescence analysis, planned two-sided Wilcoxon rank-sum tests were performed comparing HS with each tumor subtype, NOD BCC with INF BCC, and WD SCC with PD SCC. P values were adjusted within each YAP/TAZ readout using the Benjamini-Hochberg method. To assess whether stromal YAP/TAZ positivity was explained by total stromal cellularity, density-adjusted residuals were calculated by regressing logit-transformed YAP⁺ or TAZ⁺ stromal-cell fractions against log1p-transformed total stromal cell density. In addition, tumor-only linear models were fitted with YAP/TAZ readouts as outcomes and tumor entity, aggressive subtype status, and log1p-transformed total stromal cell density as predictors. For coefficient visualization, outcomes and total stromal cell density were standardized before model fitting.

For single-cell pathway and gene-set heatmaps shown in Supplementary Fig. 5C-D, PROGENy pathway activity scores and MSigDB C2/WikiPathways Hippo/YAP-associated gene-set scores were summarized as mean scores across fibroblast/CAF populations and visualized descriptively. For the YAP/TAZ-associated module analysis shown in Supplementary Fig. 5E, per-cell module scores generated with AddModuleScore were summarized as sample-level mean scores for each fibroblast/CAF population before statistical testing. Sample–fibroblast population combinations with fewer than 10 cells were excluded from sample-level testing. Differences between fibroblast/CAF populations were tested using linear models followed by Dunnett’s post-hoc comparisons, with papFIB as the reference group. Supplementary Fig. 5E shows estimated differences in sample-level module scores relative to papFIB. Significance annotations indicate Dunnett-adjusted P values: *P < 0.05, **P < 0.01, and ***P < 0.001.

For CAF niche marker comparisons, paired two-sided Wilcoxon signed-rank tests were used to compare each CAF niche class with the matched far-Outside baseline using per-sample summaries. Tests required at least four samples with non-missing values in both groups. For direct CAF niche comparisons, including iCAF versus mCAF, iCAF versus myoCAF, and mCAF versus myoCAF, paired Wilcoxon signed-rank tests were applied to per-sample differences. Effect sizes were reported as the median paired difference on the transformed marker scale. P values were adjusted using the Benjamini-Hochberg method across the corresponding set of tested cell type, marker, niche-class, or pairwise niche comparisons.

For invasive-front analyses, statistical testing was performed on sample-level zone-specific densities or sample-level marker summaries. CAF–immune associations at the invasive front were tested using Spearman rank correlations between log1p-transformed CAF subset densities and immune-cell density or immune-state endpoints. Correlations were calculated within each tumor entity and were suppressed when the minimum paired-sample requirement was not met or when either variable lacked variance. Multiple-testing correction was applied within each tumor entity across all tested CAF–immune endpoint pairs using the Benjamini-Hochberg method. Correlation heatmaps show Spearman’s rho.

Correlations between collagen-gel contraction capacity, flow-cytometry-derived percentages of MCAM⁺ and CD82⁺ cells, and qPCR-derived gene-expression readouts were assessed using two-sided Spearman rank correlations. P values from pairwise correlation tests were adjusted using the Benjamini-Hochberg method. For comparison of contraction capacity between MCAM-high and MCAM-low fibroblast cultures, cultures were stratified by a median split of the MCAM⁺ cell fraction and compared using a two-sided Wilcoxon rank-sum test. Group-wise qPCR comparisons were performed separately for each target gene using one-way ANOVA followed by Dunnett’s multiple-comparisons test, with healthy-skin fibroblasts as the reference group; comparisons were restricted to healthy-skin fibroblasts versus CAFs from each tumor type.

For functional marker heatmaps, marker intensities were summarized at the sample level before group-level visualization. Directional significance annotations were displayed on the group, zone, or niche class with the higher mean value within each planned comparison to indicate the direction of enrichment. Unless otherwise stated, data are shown as mean ± SEM. Significance was indicated as q < 0.10 and q < 0.05 for invasive-front correlation heatmaps, and as q < 0.05, q < 0.01, q < 0.001, and q < 0.0001 for CAF niche and other group-comparison analyses.

### Software

Image analysis and segmentation were performed in HALO v3.6.4134.464 using the HighPlex FL v4.2.14 plugin and HALO AI DenseNet V2 / Nuclei Seg (Indica Labs, Albuquerque, NM, USA). Single-cell data preprocessing and clustering were performed in R v4.4.0 using Seurat v5.3.0^59^ and harmony v1.2.3^60^. Pathway and module scoring of single-cell data was performed in R using decoupleR^61^, msigdbr v7.5.1, and Seurat AddModuleScore. Statistical analyses were performed in R using base R functions and multcomp v1.4-30^62^, and in GraphPad Prism v9.5.1 (GraphPad Software, Boston, MA, USA). Spatial analyses were performed in Python v3.12.4 using Squidpy v1.6.1^63^. Data visualization and statistical summaries were generated in R using standard packages, including ggplot2, pheatmap v1.0.13, ComplexHeatmap^64^, circlize^65^, and patchwork. Generative AI tools, including ChatGPT and Gemini, were used to assist with language editing and with drafting, troubleshooting, and refining R and Python scripts. All AI-assisted text, code, analyses, and outputs were reviewed, edited, and verified by the authors.

## Data availability

Source data underlying the quantitative graphs, heatmaps and statistical summaries in the main and supplementary figures are provided with this paper. Additional processed single-cell imaging mass cytometry data, spatial coordinates, cell-type annotations and spatial-analysis outputs supporting the findings of this study are available from the corresponding authors upon reasonable request, subject to institutional data-protection and ethics requirements for patient-derived tissue data and due to the large file size of the imaging datasets. The previously published single-cell RNA-sequencing dataset re-analysed in this study is available under accession number GSE254918. All other data supporting the findings of this study are available within the Article, Supplementary Information, Supplementary Tables and Source Data files.

## Code availability

Custom R and Python scripts used for image-derived data processing, statistical analysis, spatial analysis and visualization are available from the corresponding authors upon reasonable request. The analysis code will be deposited in a public repository upon acceptance of the manuscript.

## Author contribution

B.M.L. conceived the study. B.A. and B.M.L. performed and/or analyzed the majority of the experiments. B.A., J.J., N.K., I.S., D.K., and K.P. performed wet lab experiments.

B.A. performed the majority of the bioinformatic data analysis and data visualization with invaluable input and guidance from G.W., Y.Z., A.H., S.Z., A.F., and A.F.R.. P.T., K.S., E.W., A.S.C., B.G., C.F., and C.R. provided access to patient tissue. P.P. performed dermato-histopathological analysis of patient tissue. B.A. and B.M.L. wrote the manuscript with input from all authors.

## Correspondence

Correspondence and requests for materials should be addressed to Beate M. Lichtenberger and Bertram Aschenbrenner.

## Acknowledgments

We cordially thank Lena Müller from the MUW Flow Cytometry Facility for her help in IMC acquisition. Special thanks to Bärbel Reininger and Barbara Sterniczky for their technical assistance. We thank Andreas Spittler, Günther Hofbauer as well as Philipp Velicky, Sabine Rauscher and Christoph Friedl from the Flow Cytometry and Imaging core facilities of the Medical University of Vienna for their continuous support. We gratefully acknowledge Sophie Frech and Julia Deinsberger for their support with clinical data management.

This project was supported by funding granted to B.M.L. (Austrian Science Fund, FWF, V525-B28, and P36368-B; Anniversary Fund of the Austrian National Bank, OeNB, 17855; City of Vienna Fund for Innovative Interdisciplinary Cancer Research, 21059 and 22059; LEO Foundation, LF-AW_EMEA-21-400116). B.A. was financially supported by the early career seed money grant from the Austrian academy of sciences (ÖAW) and the Austrian Science Fund (FWF).

