## Supplementary Information for "Spatially organized cancer-associated fibroblast subtypes partition cutaneous carcinomas into immune-active and contracted, immune-repressed niches"

**This file contains:**

Supplementary Figures 1–5

Supplementary Table 1-5

#### Supplementary Figure 1

A

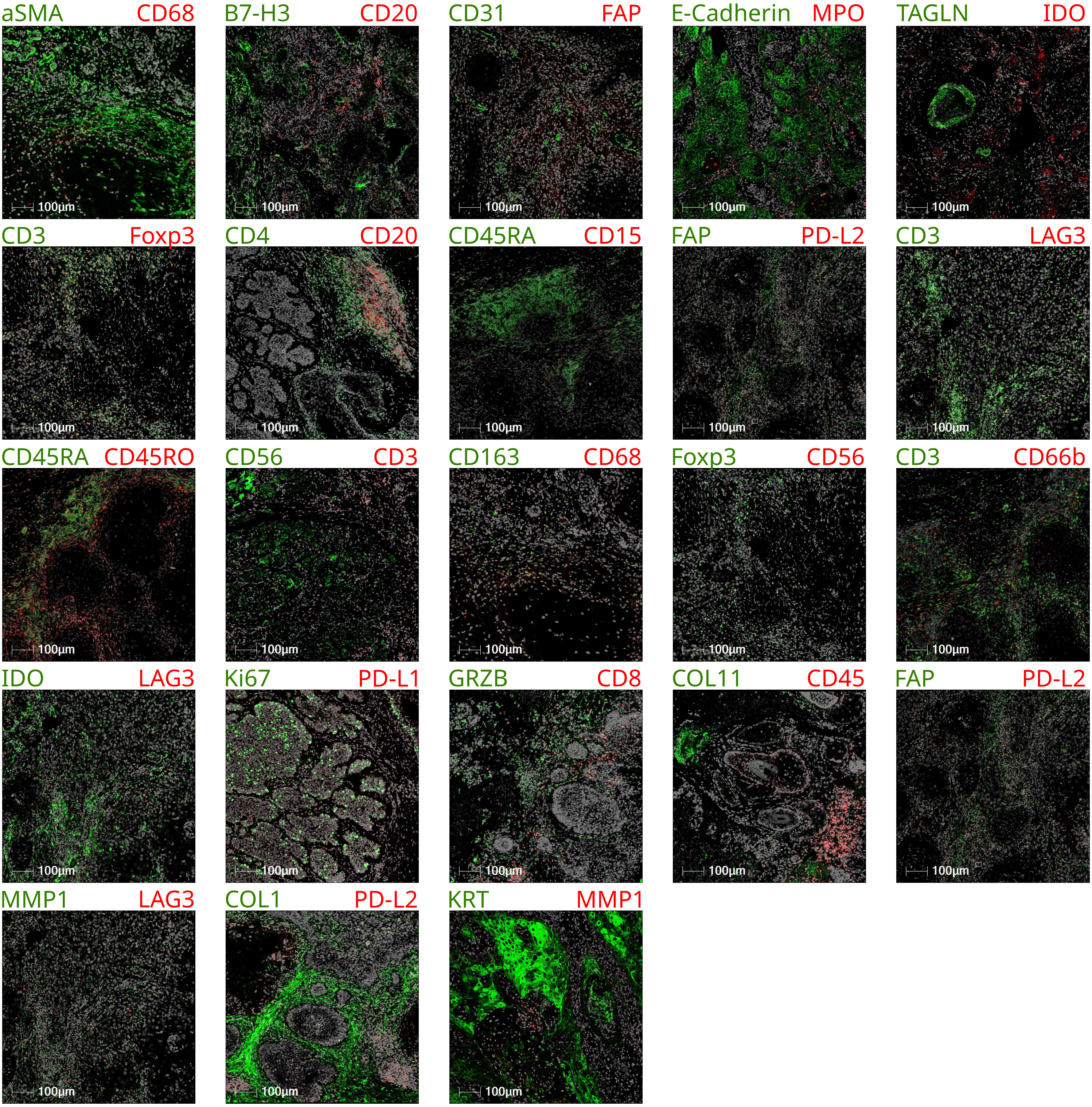


B

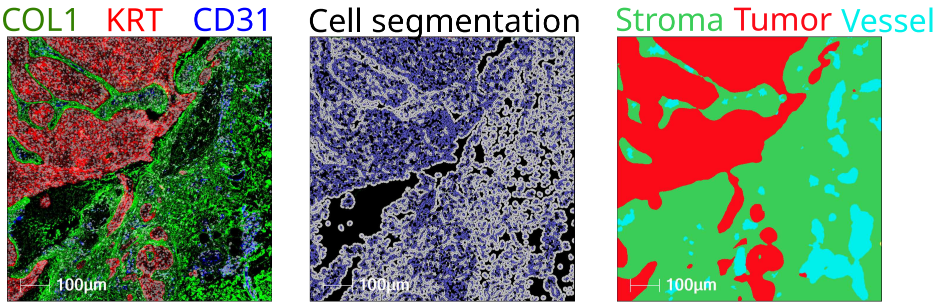

**Supplementary Figure 1. Validation of the IMC antibody panel, segmentation and tissue-classification workflow**

**(A)** Representative marker images showing expected localization and co-localization.
**(B)** Example single-cell segmentation and tissue-classification masks alongside COL1 (green), pan-KRT (red), and CD31 (blue).

#### Supplementary Figure 2

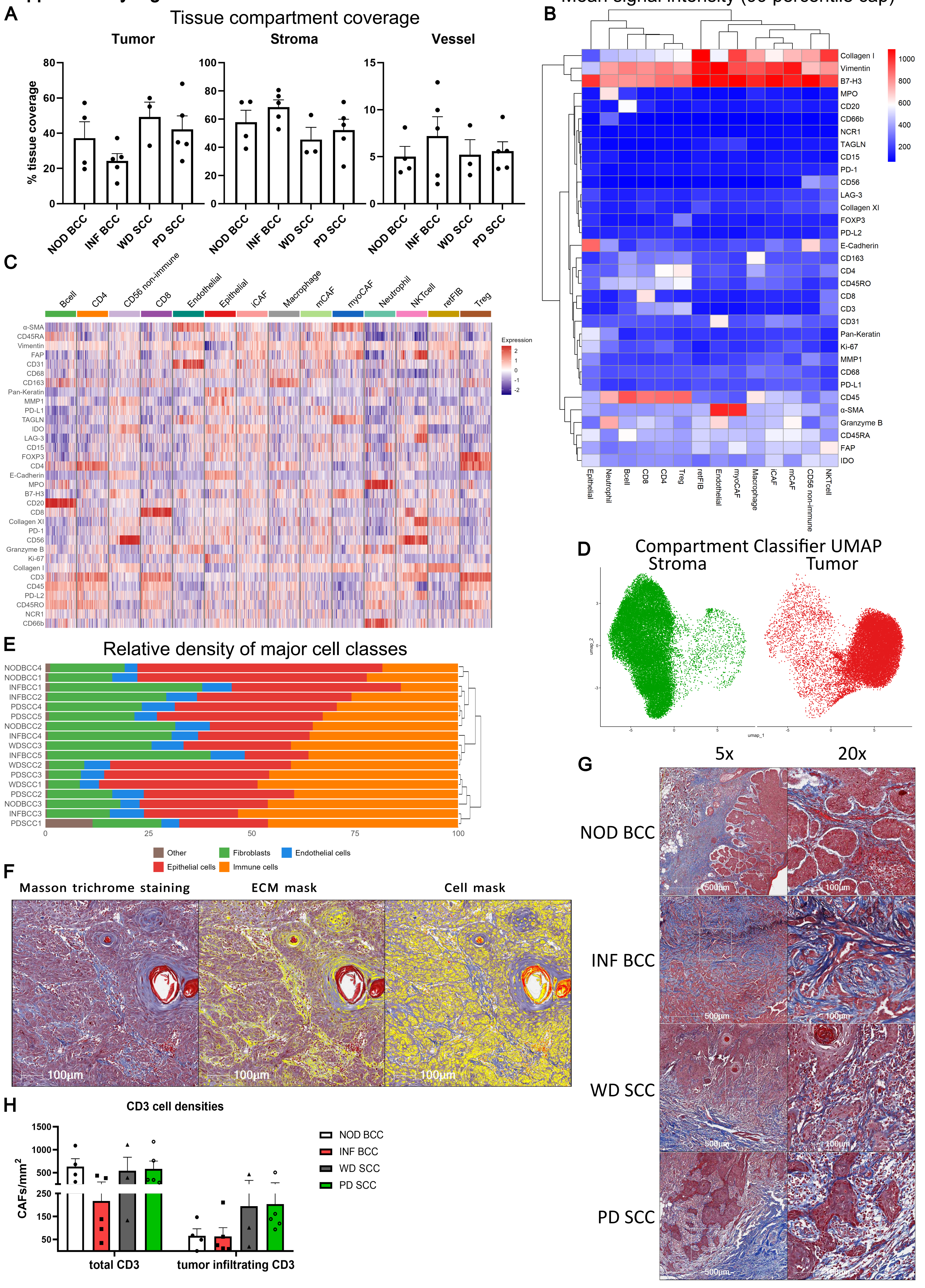

**Supplementary Figure 2. Extended tissue architecture, ECM and CD3 profiling across keratinocyte carcinoma subtypes**

**(A)** Tissue-class coverage per sample (hierarchical clustering).

**(B)** Mean marker intensity per cluster (raw values; capped at 99th percentile).

**(C)** Heatmap of representative single-cell marker expression (500 cells/cluster).

**(D)** UMAP split by tissue class (tumor vs stroma; vessels grouped with stroma).

**(E)** Per-sample relative cell-class densities.

**(F)** Masson's trichrome analysis mask generated in HALO.

**(G)** Representative Masson's trichrome images. Left: 5×, scale bar 500 µm. Right: selected region at 20×, scale bar 100 µm.

**(H)** IHC-quantified $CD3^+$ cell densities in whole tissue and within tumor nests, by tumor subtype, on sections consecutive to those used for RNA-FISH/IMC.

Supplementary Figure 3

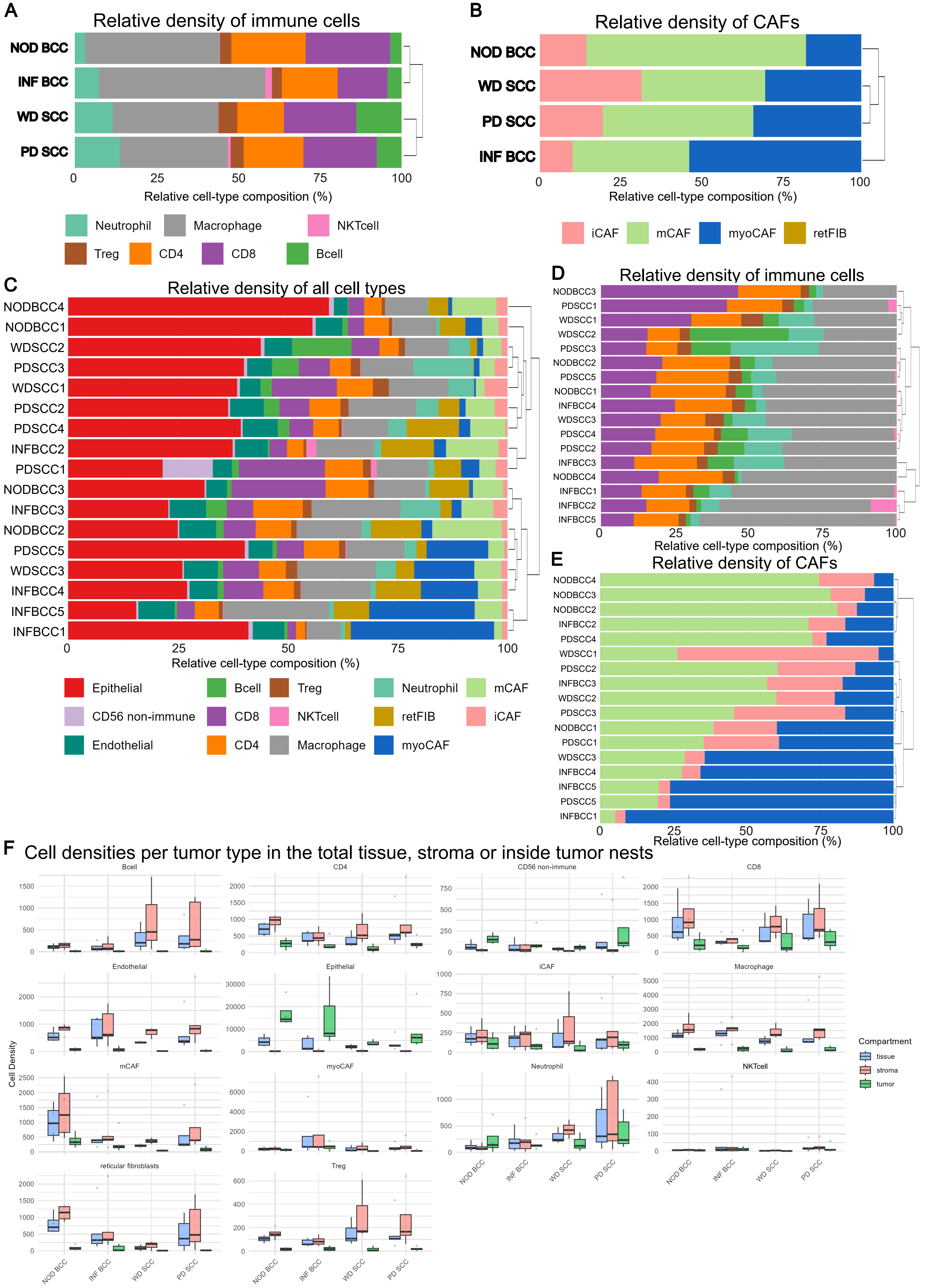

**Supplementary Figure 3. Extended cell-type composition and compartmental density analyses**

**(A)** Relative immune composition among immune cells (CD4 T, CD8 T, Treg, B, macrophage, neutrophil, NK) (%).

**(B)** Relative CAF subset composition among CAFs (iCAF, mCAF, myoCAF) (%).

**(C)** Lineage proportions per sample (% of all cells).

**(D)** Immune composition per sample (% of immune cells).

**(E)** CAF subset composition per sample (% of CAFs).

**(F)** Mean cell densities per compartment (whole tissue, stroma, tumor; cells/mm$^2$).

#### Supplementary Figure 4

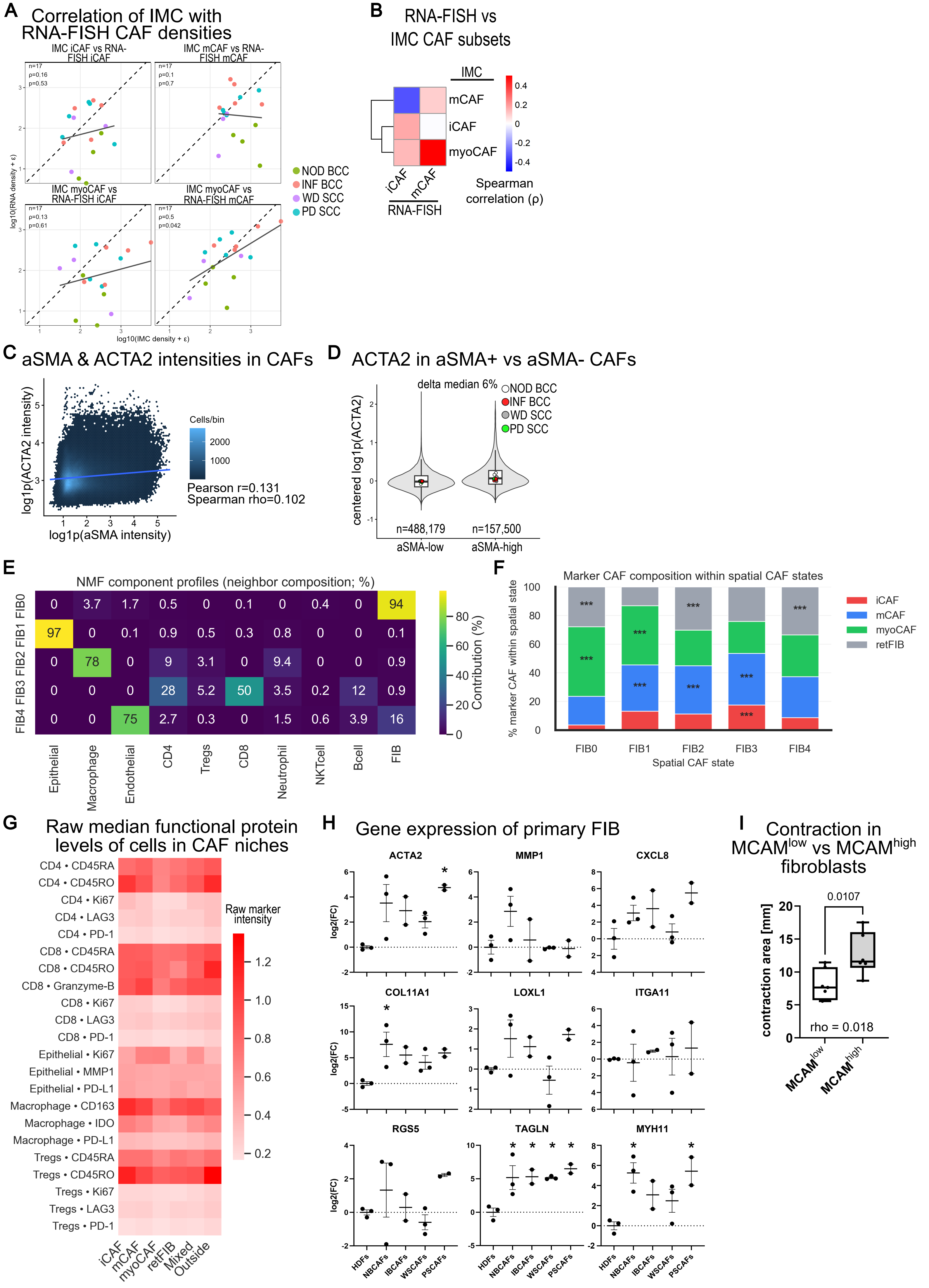

**Supplementary Figure 4. Cross-platform, spatial and functional validation of CAF programs**

**(A)** Cross-platform concordance of CAF subsets quantified by IMC and RNA-FISH: Scatter plots show per-sample CAF subset densities quantified by IMC (x-axis) and RNA-FISH (y-axis), visualized as log10(density + ε) (ε = 0.01). Points are colored by tumor type (INF BCC, NOD BCC, PD SCC, WD SCC). Dashed lines indicate the identity line (y = x). Solid lines indicate the least-squares fit across samples (shown for visualization). Spearman correlation statistics are reported per panel (n = 17 samples): IMC iCAF vs RNA iCAF, IMC mCAF vs RNA mCAF, IMC myoCAF vs RNA iCAF, and IMC myoCAF vs RNA mCAF.

**(B)** Spearman correlation (ρ) between IMC-defined and RNA-FISH-defined CAF subset densities, computed per sample.

**(C)** Hexagonal binned scatterplot of single-cell intensities for CAFs showing log1p(Opal 520 Cell Intensity (αSMA)) on the x-axis and log1p(Opal 690 Cell Intensity (ACTA2)) on the y-axis, pooled across all samples. Each hexagon summarizes the number of cells falling within that 2D bin; color indicates bin count (legend). The blue line shows the best-fit ordinary least squares regression (linear model) on the log-transformed values. Pooled correlations were computed across all included CAFs.

**(D)** Violin plots show the distribution of centered log1p(ACTA2) intensity (Opal690) for CAFs classified as αSMA− and αSMA+. Centering was performed within each sample by subtracting the sample-wise median log1p(ACTA2) across all CAFs to remove between-sample intensity offsets. Overlaid boxplots indicate median and interquartile range. Small points denote per-sample medians for each group and are colored by tumor type (NOD BCC, INF BCC, WD SCC, PD SCC). Numbers under the violins indicate total CAF counts pooled across all included samples.

**(E)** NMF component profiles showing the relative contribution (%) of each neighbor cell class to each spatial FIB state (FIB0–FIB4), derived from the NMF component matrix.

**(F)** Marker CAF composition within each spatial CAF state (stacked bars; columns sum to 100%). Asterisks indicate marker–state pairs significantly enriched by one-sided Fisher's exact test after BH-FDR correction.

**(G)** Raw median functional protein levels of cells in CAF niches (corresponding to Fig. 5G)

**(H)** Patient-level qPCR analysis of selected CAF-associated genes in primary HDFs and patient-derived CAF cultures from the indicated tumor subtypes. Each dot represents an independent patient-derived fibroblast culture.

**(I)** Primary CAF samples were stratified into Low MCAM and High MCAM groups based on the median MCAM protein levels and their contraction capacity was compared. Boxplots display the median (center line) and interquartile range (box); whiskers extend to values within 1.5× IQR. Points represent individual samples. Group differences in contraction were assessed using a two-sided Wilcoxon rank-sum test, revealing significantly higher contraction in the High MCAM group ($p = 0.018$).

### Supplementary Figure 5

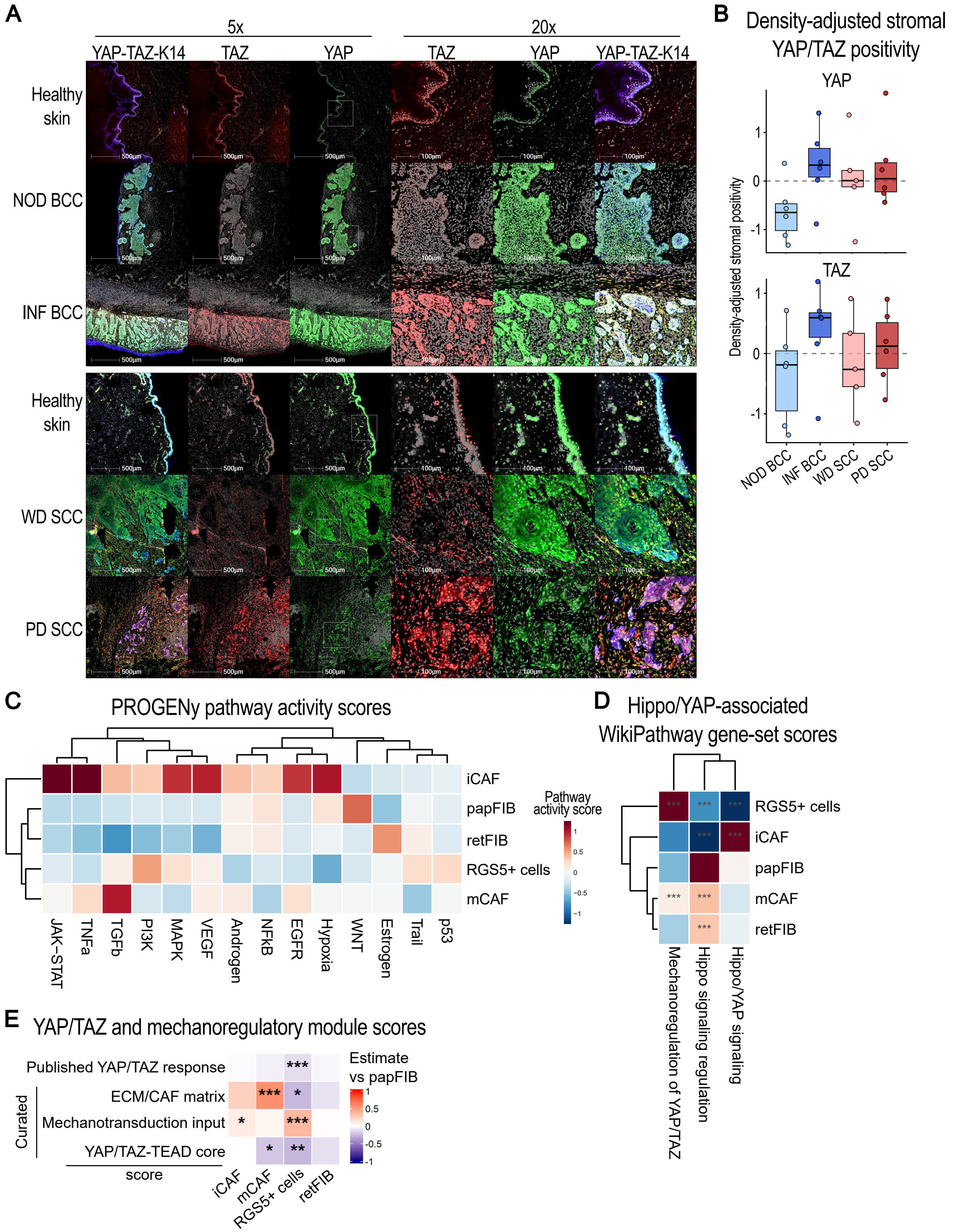

**Supplementary Figure 5. Extended stromal YAP/TAZ activation across keratinocyte carcinoma subtypes**

**(A)** Representative multiplex immunofluorescence images of YAP and TAZ staining in healthy skin, NOD BCC, INF BCC, WD SCC, and PD SCC. Rows show the indicated sample types, with 5x overview and 20x magnified views as indicated. DAPI is shown in grey, K14 in blue, TAZ in red, and YAP in green. Images illustrate nuclear YAP/TAZ localization in tumor-associated stroma and correspond to the quantification in Fig. 6K.

**(B)** Density-adjusted stromal YAP/TAZ positivity. Box plots show density-adjusted stromal YAP and TAZ positivity across NOD BCC, INF BCC, WD SCC, and PD SCC. Residuals were calculated after regressing logit-transformed $YAP^{+}$ or $TAZ^{+}$ stromal-cell fractions against total stromal cell density. Values above zero indicate higher YAP or TAZ positivity than expected based on stromal cellularity alone, whereas values below zero indicate lower positivity than expected. Points represent individual samples. Box plots show the median and interquartile range.

**(C)** Heatmap showing PROGENy pathway activity scores inferred with decoupleR across fibroblast/CAF populations from the published GSE254918 skin cancer single-cell RNA-sequencing dataset. Fibroblast subset annotations were retained from Forsthuber et al., 2024, and included iCAF, mCAF, papFIB, retFIB, and $RGS5^{+}$ cells. $RGS5^{+}$ cells correspond to the previously described mixed pericyte/myoCAF-like population. Colors indicate relative mean pathway activity scores across populations.

**(D)** Heatmap showing Hippo/YAP-associated WikiPathway gene-set scores calculated using decoupleR from selected MSigDB C2/WikiPathways gene sets. The selected gene sets represent Hippo/YAP signaling, Hippo signaling regulation, and mechanoregulation of YAP/TAZ via Hippo- and non-Hippo-dependent mechanisms. Colors indicate relative mean gene-set scores across fibroblast/CAF populations. These scores represent Hippo/YAP-associated gene-set activity and should not be interpreted as direct nuclear YAP/TAZ activity.

**(E)** Heatmap showing estimated sample-level differences in module scores across fibroblast populations relative to papFIBs. The analyzed modules included the published Cordenonsi/MSigDB YAP response signature and three curated modules capturing ECM/CAF matrix remodeling, mechanotransduction input and canonical YAP/TAZ–TEAD target-gene expression. Red indicates higher and blue indicates lower scores compared with papFIBs. Asterisks indicate Dunnett-adjusted P values from sample-level comparisons against papFIBs: $^{*}P < 0.05$, $^{**}P < 0.01$, $^{***}P < 0.001$.

#### Supplementary Table 1

##### IMC antibody panel

| Target | Metal | Clone | RRID | Concentration | Vendor | Conjugation | Catalog number |
|---|---|---|---|---|---|---|---|
| Alpha-smooth muscle actin | 141Pr | 1A4 | RRID:AB_2890139 | 1800 | Standard BioTools | Standard BioTools | # 3141017 |
| CD45RA | 142Nd | HI100 | RRID:AB_314406 | 100 | BioLegend | X8 | #304102 |
| Vimentin | 143Nd | D21H3 | RRID:AB_1069545 9 | 1000 | Standard BioTools | Standard BioTools | #3143027D |
| FAP | 144Nd | AF3715 | RRID:AB_2102369 | 250 | Bio-techne | X8 | #AF3715 |
| Anti-CD31 | 145Nd | EPR3094 | RRID:AB_3086679 | 100 | Standard BioTools | Standard BioTools | # 91H008145 |
| CD68 | 146Nd | KP1 | RRID:AB_563621 | 1500 | Standard BioTools | Standard BioTools | #91H012146 |
| CD163 | 147Sm | EDHu-1 | RRID:AB_2892115 | 80 | Standard BioTools | Standard BioTools | # 3147021 |
| Pan-keratin | 148Nd | C11 | RRID:AB_2938626 | 800 | Standard BioTools | Standard BioTools | # 3148020 |
| MMP1 | 149Sm | 41-1E5 | RRID:AB_2282006 | 50 | Millipore | X8 | #MAB3307 |
| PD-L1 | 150Nd | SP142 | RRID:AB_3106929 | 100 | Standard BioTools | Standard BioTools | # 3150033 |
| TAGLN | 151Eu | PA527463 | RRID:AB_2544939 | 50 | Thermo Fisher Scientific | X8 | #PA527463 |
| IDO | 152Sm | CI 998736 | RRID:AB_3658522 | 50 | Bio-techne | X8 | #MAB60302-100 |
| CD223/LAG-3 | 153Eu | D2G40 | RRID:AB_2811062 | 50 | Standard BioTools | Standard BioTools | # 3153028 |
| CD15 | 154Sm | QA19A61 | RRID:AB_756008 | 500 | BioLegend | X8 | #376302 |
| FoxP3 | 155Gd | 236A/E7 | RRID:AB_2910136 | 280 | Standard BioTools | Standard BioTools | # 3155016 |
| CD4 | 156Gd | EPR6855 | RRID:AB_2811051 | 150 | Standard BioTools | Standard BioTools | # 3156033 |
| E-Cadherin | 158Gd | 24 E10 | RRID:AB_2893074 | 280 | Standard BioTools | Standard BioTools | # 3158029 |
| MPO | 159Tb | Polyclonal | RRID:AB_307322 | 5000 | Abcam | X8 | # ab9535 |
| B7-H3 | 160Gd | AF1027 | RRID:AB_354546 | 100 | Bio-techne | X8 | # AF1027 |
| CD20 | 161Dy | H1 | RRID:AB_2811016 | 150 | Standard BioTools | Standard BioTools | # 3161029 |
| CD8a | 162Dy | CD8/144B | RRID:AB_2811053 | 400 | Standard BioTools | Standard BioTools | # 3162034 |
| Col11A1 | 163Dy | Polyclonal | RRID:AB_1140613 | 100 | Abcam | X8 | #ab64883 |
| PD-1 | 165Ho | EPR4877(2) | RRID:AB_3106909 | 50 | Standard BioTools | Standard BioTools | # 3165039 |
| CD56 | 166Er | 123C3 | RRID:AB_1102208 | 200 | Biorad | X8 | #MCA2693 |
| Granzyme B | 167Er | EPR20129-217 | RRID:AB_2811057 | 400 | Standard BioTools | Standard BioTools | # 3167021 |
| Ki-67 | 168Er | B56 | RRID:AB_2811061 | 100 | Standard BioTools | Standard BioTools | # 3168022 |
| Collagen 1 | 169Tm | Polyclonal | RRID:AB_2810857 | 700 | Standard BioTools | Standard BioTools | # 3169023 |
| CD3 | 170Er | Polyclonal | RRID:AB_2811048 | 150 | Standard BioTools | Standard BioTools | # 3170019 |
| CD45 | 171Yb | D3F8Q | RRID:AB_2799780 | 250 | Cell Signaling Technology | X8 | #70257 |
| PD-L2 | 172Yb | D7U8C | RRID:AB_2799999 | 100 | Standard BioTools | Standard BioTools | #3172031D |
| CD45RO | 173Yb | UCHL1 | RRID:AB_2811052 | 300 | Standard BioTools | Standard BioTools | # 3173016 |
| NCR1 | 174Yb | MM0491-8F24 | RRID:AB_3253854 | 20 | Novus | X8 | #NBP2-11820 |
| CD66b | 175Lu | G10F5 | RRID:AB_3686629 | 80 | BD Biosciences | X8 | # 571640 |
| Histone 3 | 176Yb | D1H2 | RRID:AB_2811058 | 1150 | Standard BioTools | Standard BioTools | # 3176023 |
| Nucleic acid | 191Ir 193Ir | | | 100 | Standard BioTools | Standard BioTools | #201192A |
| ISCK1 | 195Pt | Segmentation Kit | RRID:AB_3662095 | 200 | Standard BioTools | Standard BioTools | # 201500 |
| ISCK1 | 196Pt | Segmentation Kit | RRID:AB_3662095 | 200 | Standard BioTools | Standard BioTools | # 201500 |
| ISCK1 | 198Pt | Segmentation Kit | RRID:AB_3662095 | 100 | Standard BioTools | Standard BioTools | # 201500 |

#### Supplementary Table 2

##### Materials and Reagents

##### Antibodies

| Target | Catalog number | Vendor | Clone | RRID | Concentration |
|---|---|---|---|---|---|
| Mouse Anti-CD146 Monoclonal Antibody, Phycoerythrin Conjugated | # 550315 | BD Biosciences | P1H12 | RRID:AB_393604 | 10 |
| CD82 | # 564341 | BD Biosciences | 423524 | RRID:AB_2738755 | 80 |
| CD3 | # ab16669 | Abcam | SP7 | RRID:AB_443425 | 200 |
| Alexa Fluor® 488 anti-human CD324 (E-Cadherin) Antibody | #324110 | Biolegend | 67A4 | RRID:AB_756072 | 100 |
| FITC anti-human CD326 (EpCAM) Antibody | #324204 | Biolegend | 9C4 | RRID:AB_756084 | 100 |
| Rat Anti-Human CD49f Monoclonal antibody, Fitc | #MCA699F | Bio-Rad | NKI-GoH3 | RRID:AB_324269 | 20 |
| Human Fibroblast Activation Protein alpha /FAP PE-conjugated Antibody | #FAB3715P | R and D Systems | 427819 | RRID:AB_3086725 | 20 |
| Alexa Fluor(R) 647 anti-human CD90 (Thy1) | #328116 | Biolegend | Clone 5E10 | RRID:AB_893439 | 20 |
| PE/Cyanine7 anti-human CD140a (PDGFRα) | #323508 | Biolegend | Clone 16A1 | RRID:AB_2565597 | 20 |
| PE anti-human Podoplanin | #337004 | Biolegend | Clone NC-08 | RRID:AB_1595457 | 100 |
| YAP | #14074 | Cell Signaling Technology | D8H1X | RRID:AB_2650491 | 100 |
| TAZ/WWTR1 | #AMAB90730 | Merck | CL0371 | | 50 |
| Keratin 14 | #906004 | Biolegend | Poly9060 | RRID:AB_2616962 | 250 |
| goat anti-rabbit IgG (H+L) Cross-Adsorbed Secondary Antibody, Alexa Fluor 594 | A-11012 | Thermo Fisher Scientific | | RRID:AB_2534079 | 500 |
| goat anti-mouse IgG (H+L) Cross-Adsorbed Secondary Antibody, Alexa Fluor 647 | A-21235 | Thermo Fisher Scientific | | RRID:AB_2535804 | 500 |
| goat anti-chicken IgY (H+L) cross-adsorbed Alexa Fluor 488 | A-11039 | Thermo Fisher Scientific | | RRID:AB_2534096 | 500 |

##### qPCR probes

| Target | Catalognumber | Vendor |
|---|---|---|
| MMP1 | Hs00899659_g1 | Thermo Fisher Scientific |
| CXCL8 | Hs00174103_m1 | Thermo Fisher Scientific |
| LOXL1 | Hs00935937_m1 | Thermo Fisher Scientific |
| ITGA11 | Hs01012939 | Thermo Fisher Scientific |
| COL11A1 | Hs01097664_m1 | Thermo Fisher Scientific |
| ACTA2 | Hs00426835_g1 | Thermo Fisher Scientific |
| MYH11 | Hs00975796_m1 | Thermo Fisher Scientific |
| RGS5 | Hs01591223_s1 | Thermo Fisher Scientific |
| TAGLN | Hs01038777_g1 | Thermo Fisher Scientific |
| GAPDH | Hs99999905 | Thermo Fisher Scientific |

##### qPCR reagents

| Product | Catalognumber | Vendor |
|---|---|---|
| Qiagen RNeasy Mini Kit | #74106 | Qiagen |
| RevertAid H Minus First Strand cDNA Syn | #K1631 | Thermo Fisher Scientific |
| DNaseI | #EN0521 | Thermo Fisher Scientific |
| Taqman 2xUniversal PCR Master Mix | #4324018 | Applied Biosystems |

##### RNAScope

| Product | Catalognumber | Vendor |
|---|---|---|
| RNAscope® Probe - Hs-MMP1 | #412641-C1 | Advanced Cell Diagnostics, Bio-Techne |
| RNAscope® Probe - Hs-COL1A1-C4 | #401891-C4 | Advanced Cell Diagnostics, Bio-Techne |
| RNAscope® Probe - Hs-COL11A1-C3 | #400741-C3 | Advanced Cell Diagnostics, Bio-Techne |
| RNAscope® Probe - Hs-COL11A1-C2 | #400741-C2 | Advanced Cell Diagnostics, Bio-Techne |
| RNAscope® Probe - Hs-RGS5-C3 | #533421-C3 | Advanced Cell Diagnostics, Bio-Techne |
| RNAscope® Probe - Hs-ACTA2-O1-C2 | #444771-C2 | Advanced Cell Diagnostics, Bio-Techne |
| Multiplex Fluorescent Reagent Kit v2 | #323135 | Advanced Cell Diagnostics, Bio-Techne |
| RNAscope® 4-Plex Ancillary kit | #323120 | Advanced Cell Diagnostics, Bio-Techne |
| Opal570 | FP1488001KT | Akoya |
| Opal620 | FP1495001KT | Akoya |
| Opal690 | FP1497001KT | Akoya |
| Opal780 | FP1501001KT | Akoya |

##### Cell culture

| Product | Catalognumber | Vendor |
|---|---|---|
| Matrigel | 356231 | Corning |
| Rat-tail Collagen1 | 354236 | Corning |
| StemPro™ Accutase™ Cell Dissociation F | A1110501 | Thermo Fisher Scientific |
| Trypsin | 25200072 | Thermo Fisher Scientific |
| DMEM | 11995073 | Gibco |
| TGFB | 100-21-10ug | PeproTech® |
| FGF | 100-18B | PeproTech® |
| EGF | 234-FSE | R&D Systems |
| Insulin | I979278-5ML | Sigma-Aldrich |
| 2-phosphate ascorbic acid | SC-228390 | Santa Cruz Biotechnology |
| Hydrocortisone Hemisuccinate | H0888-1G | Sigma-Aldrich |
| Pen-Strep | 15140-122 | Gibco |

| | | |
|---|---|---|
| Gentamycin | 15710049 | Gibco |
| Normicin | ant-nr-05 | InvivoGen |
| Non-essential amino acids | 11140-050 | Gibco |
| Glutamine | 25030-024 | Gibco |

**Metal-konjugation kits**

| Product | Catalognumber | Vendor |
|---|---|---|
| Maxpar® X8 Antibody Labeling Kit, 142Nd | #201142A | Standard BioTools |
| Maxpar® X8 Antibody Labeling Kit, 144Nd | #201144A | Standard BioTools |
| Maxpar® X8 Antibody Labeling Kit, 149Sm | #201149A | Standard BioTools |
| Maxpar® X8 Antibody Labeling Kit, 151Eu- | #201151A | Standard BioTools |
| Maxpar® X8 Antibody Labeling Kit, 152Sm | #201152A | Standard BioTools |
| Maxpar® X8 Antibody Labeling Kit, 154Sm | #201154A | Standard BioTools |
| Maxpar® X8 Antibody Labeling Kit, 159Tb- | #201159A | Standard BioTools |
| Maxpar® X8 Antibody Labeling Kit, 160Gd | #201160A | Standard BioTools |
| Maxpar® X8 Antibody Labeling Kit, 163Gd | #201163A | Standard BioTools |
| Maxpar® X8 Antibody Labeling Kit, 166Er- | #201166A | Standard BioTools |
| Maxpar® X8 Antibody Labeling Kit, 171Yb- | #201171A | Standard BioTools |
| Maxpar® X8 Antibody Labeling Kit, 175Lu- | #201175A | Standard BioTools |
| Maxpar® X8 Antibody Labeling Kit, 176Yb- | #201176A | Standard BioTools |

**Immunohistochemistry**

| Product | Catalognumber | Vendor |
|---|---|---|
| Biotinylated goat anti-rabbit antibody | #BA-1000 | Vector |
| Novocastra Streptavidin-HRP | #RE7104 | Leica Biosystems |
| Dako AEC+ High sensitivity substrate | #K3469 | Dako |

### Supplementary Table 3

#### Patient information corresponding to primary fibroblasts

| Cell line | Sample type | Sex | Age | Localization |
|---|---|---|---|---|
| HDF1 | HS | f | 37 | upper arm |
| HDF2 | HS | f | 33 | upper arm |
| HDF3 | HS | f | 52 | abdomen |
| IBCAF1 | INF BCC | m | 77 | shoulder |
| IBCAF2 | INF BCC | m | 73 | cheek |
| NBCAF1 | NOD BCC | m | 80 | neck |
| NBCAF2 | NOD BCC | m | 88 | clavicular |
| NBCAF3 | NOD BCC | m | 88 | lower leg |
| PSCAF1 | PD SCC | m | 80 | head |
| PSCAF2 | PD SCC | m | 87 | head |
| WSCAF1 | WD SCC | m | 85 | lower leg |
| WSCAF2 | WD SCC | m | 81 | lower leg |
| WSCAF3 | WD SCC | m | 78 | lower leg |

### Supplementary Table 4

#### Number and types of samples spatially analyzed

| YAP/TAZ K14 staining | n=28 |
|---|---|
| HS | 7 |
| NOD BCC | 6 |
| INF BCC | 6 |
| WD SCC | 5 |
| PD SCC | 6 |

| IMC & Maason trichrome & CD3 IHC & RNA-FISH stainings | n=17 |
|---|---|
| NOD BCC | 4 |
| INF BCC | 5 |
| WD SCC | 3 |
| PD SCC | 5 |

| RNA-FISH + aSMA staining | n=12 |
|---|---|
| NOD BCC | 3 |
| INF BCC | 3 |
| WD SCC | 3 |
| PD SCC | 3 |

| Total nr. of analyzed samples | n=57 |
|---|---|
| Nr of patients analyzed | 44 |
| HS | 7 |
| NOD BCC | 13 |
| INF BCC | 14 |
| WD SCC | 11 |
| PD SCC | 14 |

#### Supplementary Table 5

##### Number and types of samples spatially analyzed

| Gene number | CORDENONSI YAP (MSigDB C6) | YAP/TAZ-TEAD core (curated) | Mechano-input (curated) | ECM/CAF matrix (curated) |
|---|---|---|---|---|
| 1 | AGFG2 | CYR61 | ITGA1 | COL1A1 |
| 2 | AMOTL2 | CTGF | ITGA2 | COL1A2 |
| 3 | ANKRD1 | ANKRD1 | ITGA5 | COL3A1 |
| 4 | ASAP1 | AMOTL2 | ITGAV | COL5A1 |
| 5 | AXL | AXL | ITGA11 | COL5A2 |
| 6 | BICC1 | SERPINE1 | ITGB1 | COL6A1 |
| 7 | BIRC5 | THBS1 | ITGB3 | COL6A2 |
| 8 | CYR61 | NT5E | PTK2 | COL6A3 |
| 9 | CTGF | F3 | SRC | COL11A1 |
| 10 | CDC20 | NUAK2 | TLN1 | COL12A1 |
| 11 | CDKN2C | LATS2 | TLN2 | FN1 |
| 12 | CENPF | PTPN14 | VCL | POSTN |
| 13 | COL4A3 | | PXN | TNC |
| 14 | CRIM1 | | ZYX | THBS1 |
| 15 | DAB2 | | RHOA | SPARC |
| 16 | DDAH1 | | RHOC | VCAN |
| 17 | DLC1 | | ROCK1 | LOX |
| 18 | DUSP1 | | ROCK2 | LOXL1 |
| 19 | DUT | | DIAPH1 | LOXL2 |
| 20 | ECT2 | | DIAPH3 | MMP2 |
| 21 | EMP2 | | MYH9 | MMP11 |
| 22 | ETV5 | | MYH10 | MMP14 |
| 23 | FGF2 | | MYL9 | TIMP1 |
| 24 | FLNA | | MYL12A | TIMP2 |
| 25 | FSCN1 | | MYL12B | ITGA11 |
| 26 | FSTL1 | | ACTA2 | BGN |
| 27 | GADD45B | | TAGLN | FBN1 |
| 28 | GAS6 | | CNN1 | PLOD2 |
| 29 | GGH | | ACTN1 | |
| 30 | GLS | | ACTN4 | |
| 31 | HEXB | | FLNA | |
| 32 | HMMR | | FLNB | |
| 33 | ITGB2 | | CAV1 | |
| 34 | ITGB5 | | ILK | |
| 35 | MARCKS | | FERMT2 | |
| 36 | MDFIC | | TNS1 | |
| 37 | NDRG1 | | LIMS1 | |
| 38 | PDLIM2 | | PARVA | |
| 39 | PHGDH | | PARVB | |
| 40 | PMP22 | | | |
| 41 | SCHIP1 | | | |
| 42 | SERPINE1 | | | |
| 43 | SGK1 | | | |
| 44 | SH2D4A | | | |
| 45 | SHCBP1 | | | |
| 46 | SLIT2 | | | |
| 47 | STMN1 | | | |
| 48 | TGFB2 | | | |
| 49 | TGM2 | | | |
| 50 | THBS1 | | | |
| 51 | TK1 | | | |
| 52 | TNNT2 | | | |
| 53 | TNS1 | | | |
| 54 | TOP2A | | | |
| 55 | TSPAN3 | | | |
